# Mushrooms arising from the mole latrine reveal the life of talpid moles: proposals of ‘myco-talpology’ and ‘habitat-cleaning symbiosis’

**DOI:** 10.1101/2022.10.22.513302

**Authors:** Naohiko Sagara, Nobuko Tuno, Yu Fukasawa, Shin-ichiro Kawada, Taiga Kasuya

## Abstract

Myco-talpology is the science of moles and mushrooms based on our discovery in 1976 that underground nests of talpid moles can be located by the aboveground fruiting of the agarics *Hebeloma danicum* or *Hebeloma sagarae*, a Japanese sibling species of European *Hebeloma radicosum*. Hyphae of these mushrooms specifically colonise mole latrines near nests, forming ectomycorrhizas with the roots of their host trees. By this process, the hyphae and roots absorb, transform, and translocate the nutrients from mole excretions, cleaning the mole’s habitat. The mushrooms fruit on the ground with thick root-like tissue stretching up from the ectomycorrhizas. The presence of the fruit bodies serves as the only indicator for underground mole nests, enabling us to provide novel insights into mole ecology. We excavated 74 *H. sagarae* or *H. danicum* sites in Japan and 13 *H. radicosum* sites in Europe and identified the nesting species. These investigations provided new perspectives on latrine making, nest structure, breeding and nestlings, the neighbouring presence of different Talpini species, and long-term nesting at the same site with inhabitant changes. Moreover, they suggested that a tripartite habitat-cleaning symbiosis among moles, mushrooms, and mushroom-host trees in the latrine must have existed for a long time in Japanese and European forests. While moles are traditionally considered inhabitants of open areas, their special relationship with mushrooms and trees suggests they may originally be forest inhabitants.

## 1. Introduction

The ecology of the family Talpidae (talpid moles: Soricomorpha, Mammalia) appears poorly understood because of their subterranean life habits, particularly their nesting. The daily lives of moles are centred on their nests (Haeck, 1969 in Gorman and Stone [1990], Gorman and Stone, 1990). Therefore, research on the ecology of moles should begin with locating their nests. In open land with shallower soil or higher water level in Europe, *Talpa europaea* L. construct nests in huge mounds easily recognised as ‘fortresses’ (Godfrey and Crowcroft, 1960; Gorman and Stone, 1990). Similarly, in North America, *Scapanus townsendii* Bachman construct recognisable ‘nest mounds’ above breeding nests (Kuhn et al., 1966). However, these discernible cases are exceptional; most nests remain undiscovered by field surveys. Particularly in the forest (defined as a tree-covered area), talpid mole nests have no aboveground signs of their presence. For this reason and a lack of human interest, the lives of moles in the forest have been poorly documented and moles’ ecological information has been limited mostly to that from trapping or tracking in open land.

In this context, Sagara (1978, 1980) found that two agaric mushroom species occur (fruit) independently or together on mole nests, colonising nearby deserted or old latrines. The occurrence of these mushrooms is the only clue for locating mole nests in the forest without using scientific instrumentation such as radioisotope tracing or radio tracking. This observation has led to the exploration of the life habits of forest-dwelling moles based on their nests, an approach tentatively termed ‘myco-talpology’ (myco-= mushrooms and talpo-= talpids) that can illuminate the natural history of talpid moles. Based on myco-talpology, we first clarified the range of mammal species involved and other conditions associated with this phenomenon and then investigated questions that arose. There have been no similar attempts from other countries than Japan.

This myco-talpology approach identified unknown aspects of mole life. In particular, the mole-mushroom-tree relationship should be considered a ‘habitat-cleaning symbiosis’ that may be essential, in which mushroom hyphae and tree roots colonise the mole latrine, forming mycorrhizas, and clean it. Indeed, the term ‘mole latrine’ has not been used in the mole ecology literature.

Myco-talpology has also inevitably led to the perspective that mammals other than moles may replace them in the tripartite association. However, most observed cases concern moles (**Table 1**), and the primary findings of this study relate to moles. Therefore, we consider the term myco-talpology to have continued relevance, and this article summarises our understanding of myco-talpology since 1976.

**Table 1.**
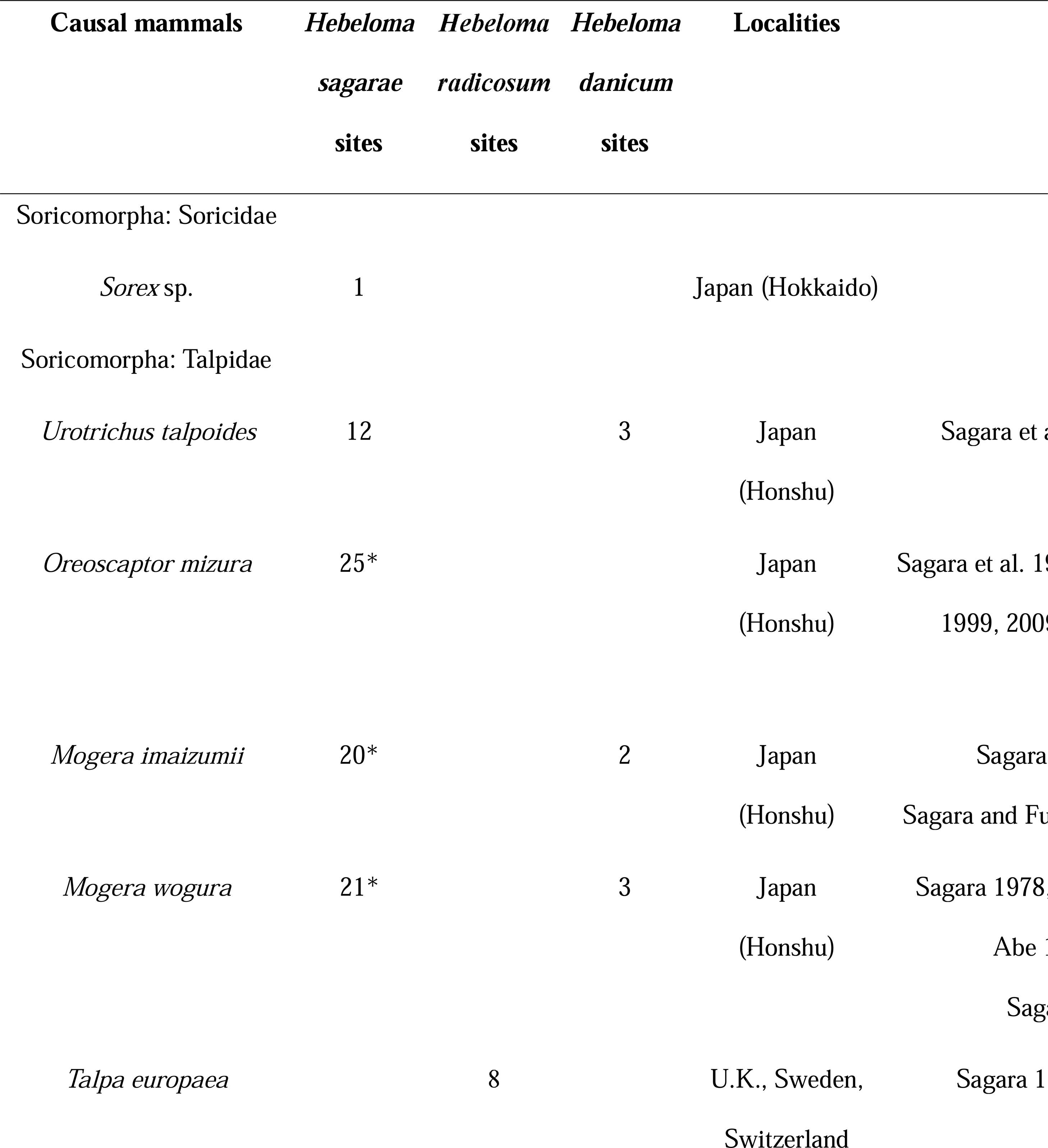
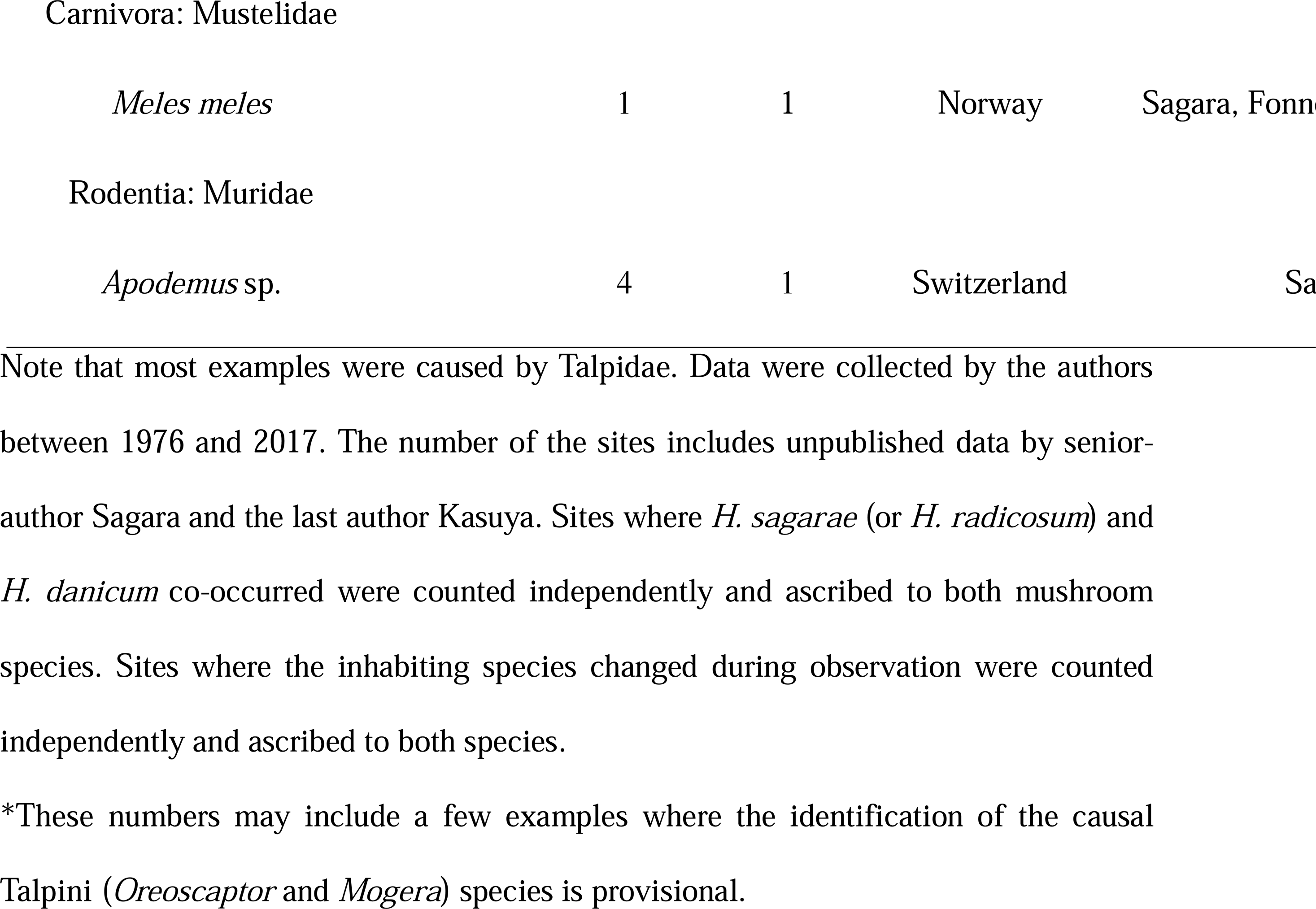
Mammals whose nesting causes the occurrence of mushrooms *H. sagarae*, *H*. *radicosum*, or *H. danicum*, and the number of sites where this mammal-mushroom association was observed.

## 2. General methods

### 2.1 Organisms and study area

The mushrooms involved are shown in **Figure 1**. Eberhardt et al. (2020) recently proposed, following Kasuya et al. (2013), one of these two species, *Hebeloma sagarae* T. Kasuya, Mikami, Beker, and U. Eberh., as a Japanese species distinct from European *Hebeloma radicosum* (Bull.) Ricken. They were jointly named *H. radicosum* in our previous publications. *Hebeloma danicum* Gröger denotes *Hebeloma spoliatum* sensu Lange (1938) and sensu Hongo (1987) as discussed by Sagara et al. (2000, 2008c), which was reported as *H. spoliatum* in our publications prior to 2000. Al these species are ectomycorrhizal (**Sections 3.1** and **3.3**).

**Figure 1.**
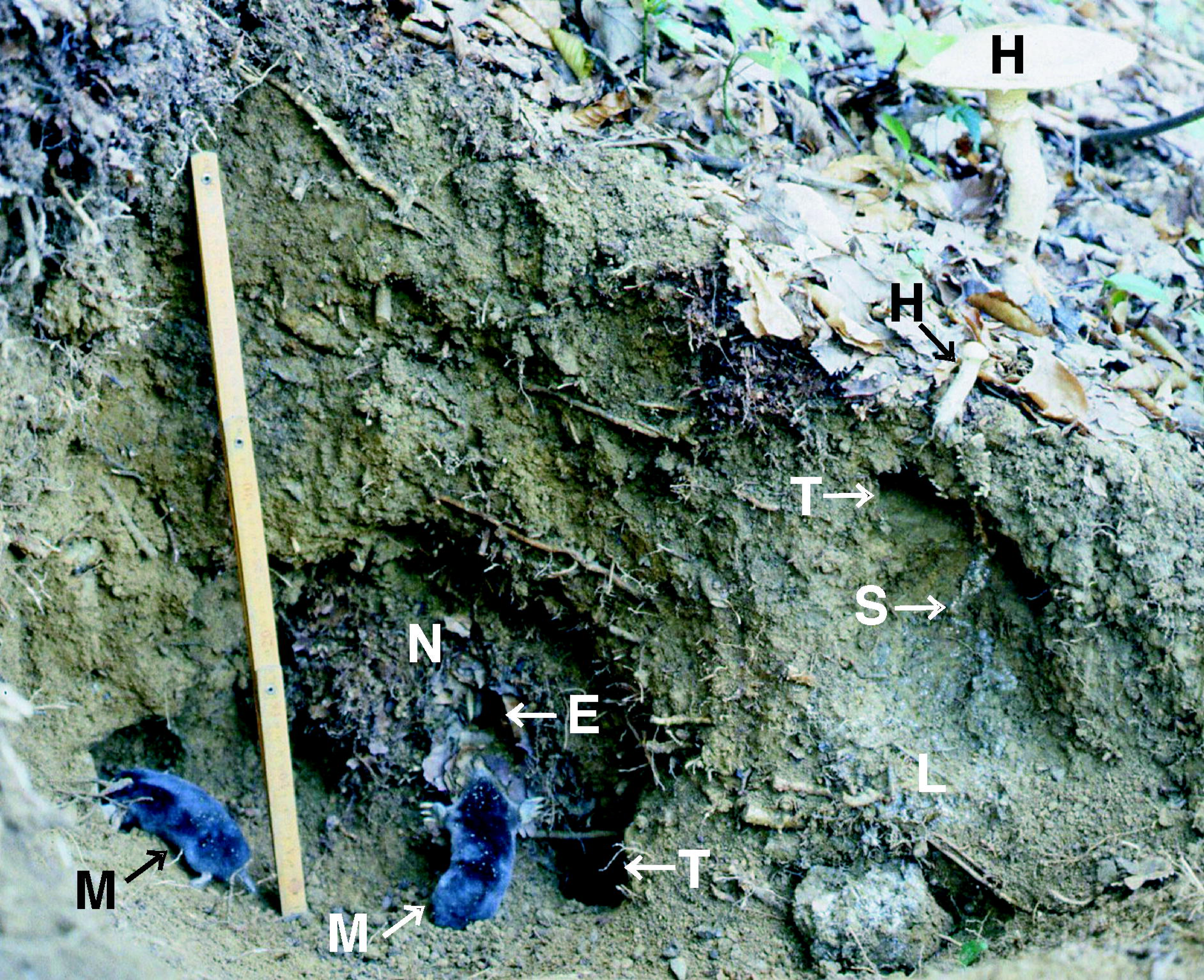
Mushrooms involved. (**A**, **B**) *H. sagarae*, (**C**) *H. danicum*. In Europe, *H. sagarae* is replaced by *H. radicosum.* Each fruit body has a rooting stipe from the mole’s deserted latrine. Arrows indicate ground level. The fruit bodies in **A** lack the lower (basal) parts of their stipes, indicating they originated deeper in the soil. Bar denotes 10 cm (for A-C). From Sagara (1998b), reproduced with permission.

The animals involved were, in most cases, talpid moles (Sagara, 1999; **Table 1**, **Figure 2**). Our study was performed primarily in central Japan (Kyoto Prefecture and adjacent districts), where four talpid species are common and often overlap in their distribution: *Urotrichus talpoides* Temminck (greater Japanese shrew-mole), *Oreoscaptor mizura* (Günther) (Japanese mountain mole), *Mogera imaizumii* (Kuroda) (lesser Japanese mole), and *Mogera wogura* (Temminck) (large Japanese mole). An area with such a diverse mole distribution is rare, and all these species are associated with the mushrooms in **Figure 1**.

**Figure 2.**
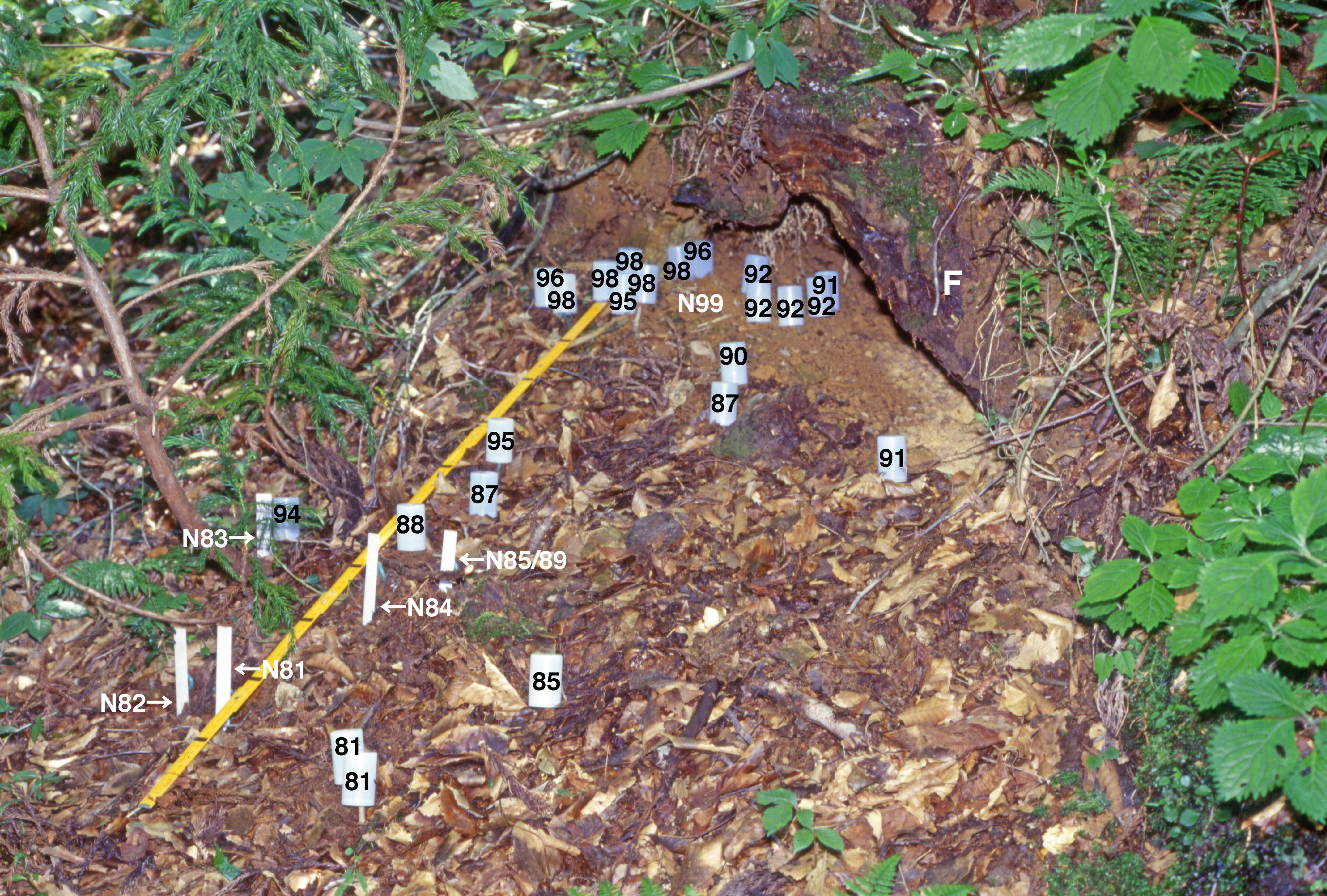
Talpid moles that occurred in central Japan and were treated in this study as causal animals for the occurrence of mushrooms *H. sagarae* and *H. danicum*: (**A**) *U. talpoides*; (**B**) *O. mizura*; (**C**) *M. imaizumii*; (**D**) *M. wogura*. **A** from Daimonji-yama, Kyoto City (col. O. Murakami), **B**-**D** from Ashiu, Miyama-cho, Nantan-shi, Kyoto Prefecture (col. N. Sagara). The nails of *O. mizura* stretched out abnormally during 55 days of captivity. Bars denote 10 cm. Photographs: **A** original, **B-D** from Sagara et al. (1989), reproduced with permission.

This study extended to some regions of Europe, where the European mole *T. europaea* is found (**Figure 3**), leading to the discovery that the wood mouse *Apodemus* may replace moles in the animal-fungus-plant relationship (**Table 1**). Furthermore, we have paid special attention to some northern areas devoid of talpids but with those mushrooms: Ireland and Norway in Europe and Hokkaido in Japan. As a result, we found that the badger *Meles meles* L. in Norway and the shrew *Sorex* sp. in Hokkaido replaced the moles (**Table 1**).

**Figure 3.**
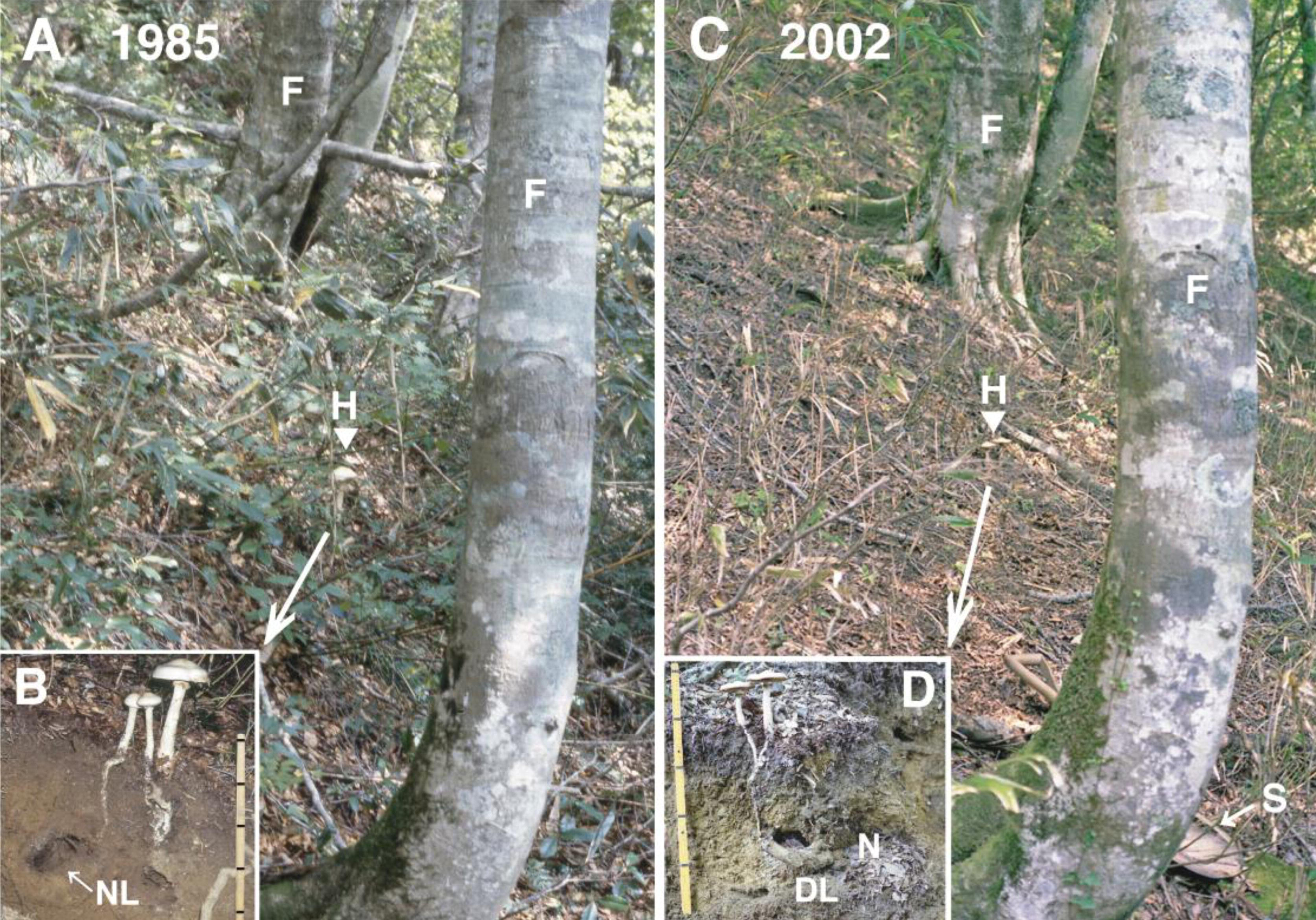
An excavation in Europe to study the relationship between mushrooms and moles (Sagara, unpublished). (**A**) Site of the mushroom *H. radicosum* occurrence at Farner, Oberlangenegg, Bern, Switzerland, 915 m alt. Note the presence of beechs at the particular site while the area was dominated by conifers *Picea abies* and *Abies alba*. Key: H, the point at which the mushroom fruit bodies were found and marked on 29 September 1997; F, beech *F. sylvatica* trees with their leaves scattered on the ground. (**B**) Excavation at this site on 25 November 1997. Key: P, pickaxe; S, shovel. (**C**) Mole nest (N) is located beneath the trunk of a *P. abies* tree. Key: E, entrance to the resting cavity within the nest. Two thick lines on the folding scale are at a 10-cm interval. (**D**) *T. europaea* nest removed from this location. Key: E, the same entrance as shown in **C**. Photographs: **A**, **C** original; **B**, **D** by Guido Bieri, reproduced with permission.

The nomenclature for mammals follows Ohdachi et al. (2009) for Japanese species except *O. mizura* and Wilson and Reeder (2005) for European and North American species. *O. mizura* was reclassified by Kawada (2016) and had been treated as *Euroscaptor mizura* in our previous papers. *M. wogura* and *Mogera kobeae* in the articles by Sagara et al. before 1998 have been renamed *M. imaizumii* and *M. wogura*, respectively, in subsequent papers based on the revision by Motokawa and Abe (1996).

The plants involved were ectomycorrhizal broad-leaved trees (Sagara, 1999; **Table 2**). Consequently, the study area extended to mountains and hills vegetated by those trees. Vegetation of the studied sites was recorded by the names of trees dominating each site, primarily as an environment but also as the forests containing possible mycorrhizal hosts for those mushrooms. The authors of the scientific tree names are omitted.

**Table 2.**
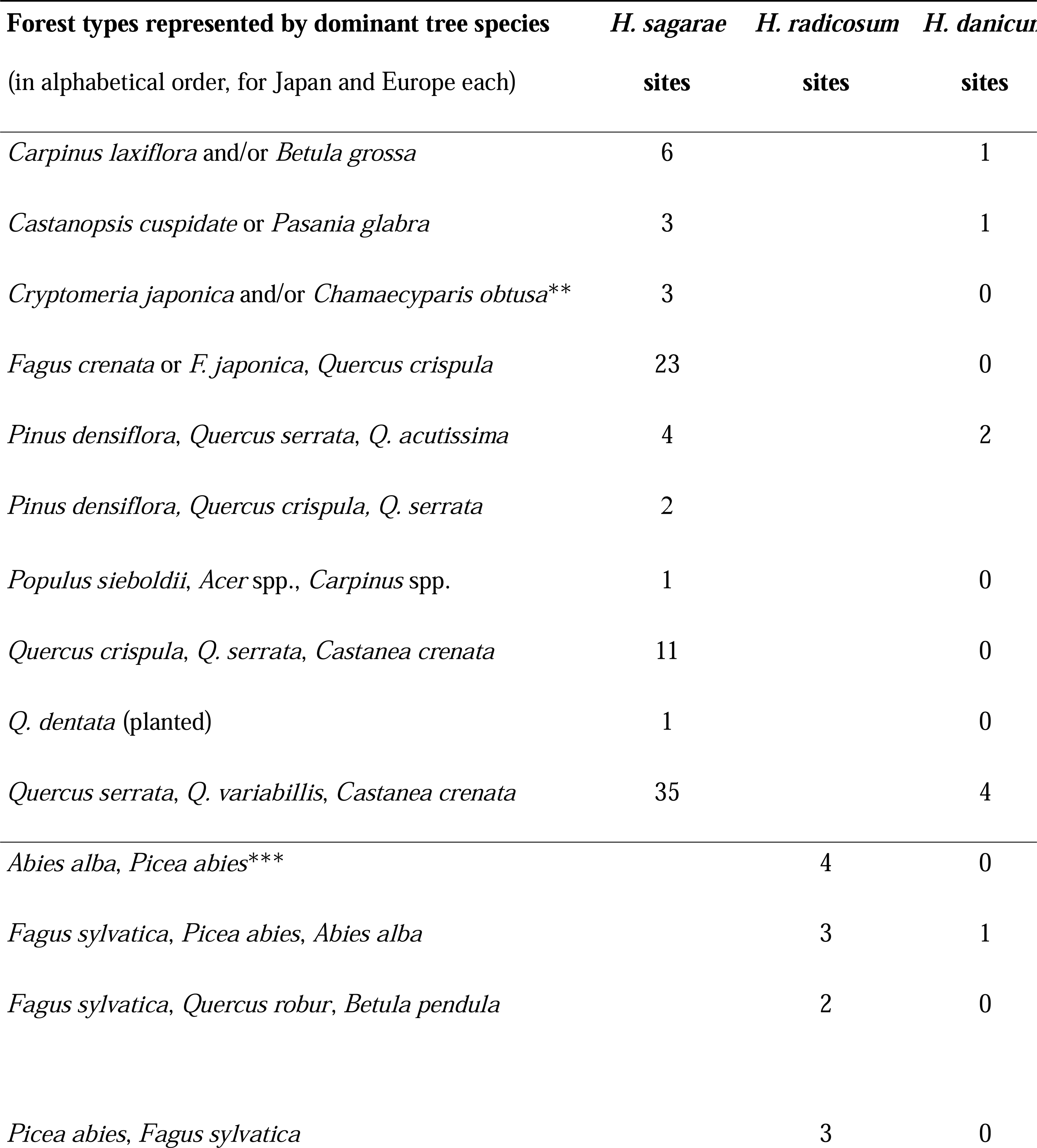
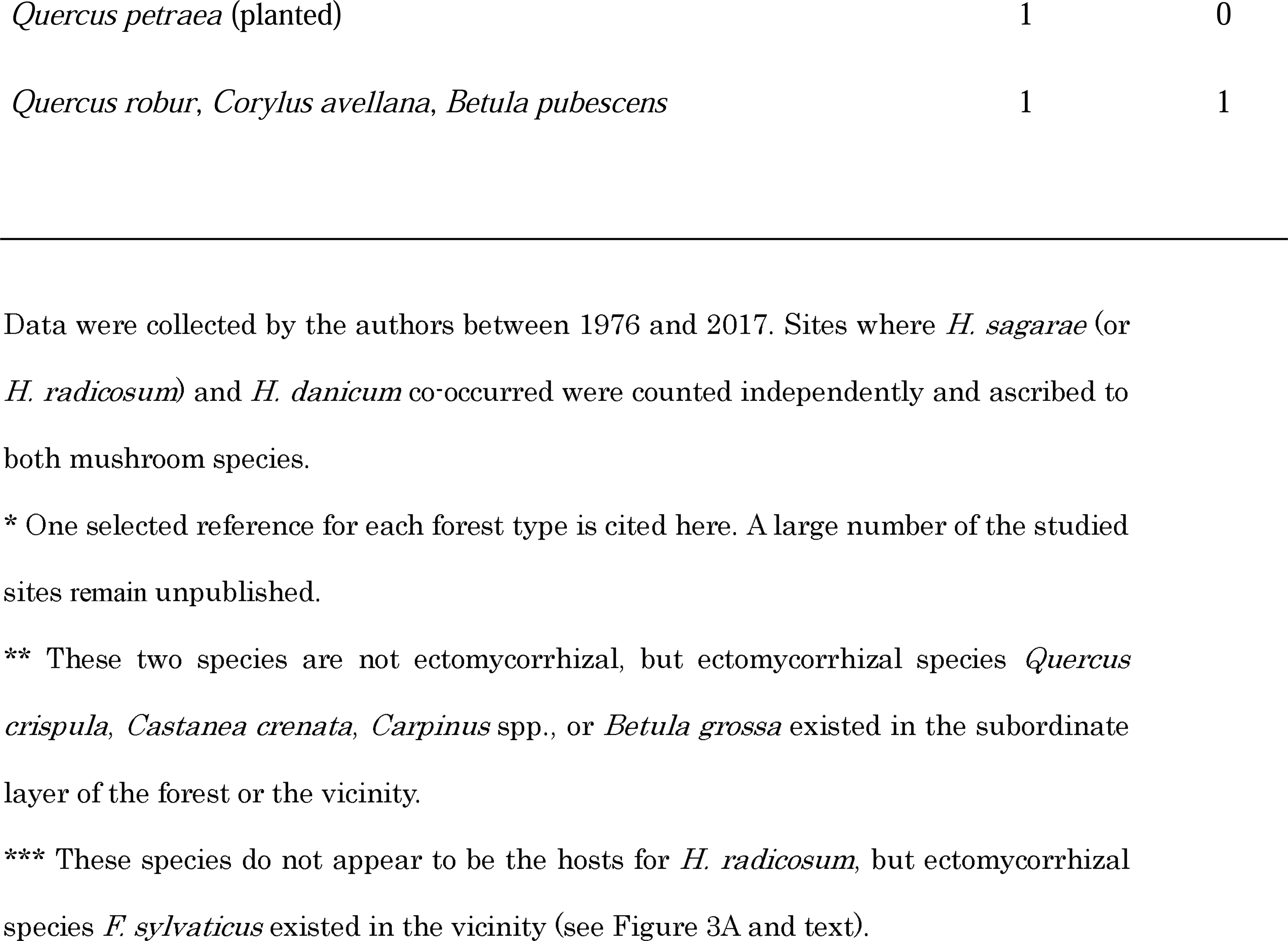
Forest types and site numbers therein of the occurrence of mushrooms Hebeloma sagarae, H. radicosum, or H. danicum, showing their association with putative ectomycorrhizal trees of Fagaceae, Betulaceae, or Salicaceae.

### 2.2 Locating mushroom occurring sites

When the occurrence (fruiting) of *H. sagarae*, *H. danicum*, or *H. radicosum* was observed, the points of the fruit bodies were marked with pegs for later studies (cf. H in **Figure 3A**). These mushrooms were rarely brought into our study: senior-author Sagara studied 86 mushroom sites, including 13 in Europe, over 38 years from 1976 to 2014, averaging only 2.5 finds per year (**Table 1**). Furthermore, among those 86 sites, he found only 12 (12.5%) by himself; local mycologists found and told him about the others. The cooperation and assistance among mycologists have been essential.

We do not report the number of mushroom fruit bodies found at each site since they are neither animals nor plants (Whittaker, 1969). Mushroom fruit bodies are not biological ‘individuals’ as recognised in animals and plants, and it is almost meaningless to consider how many fruit bodies exist per nest.

Mushroom fruiting appears to occur even one year later with disused latrines, as mycorrhizal colonisation may occur or continue after disuse. How long fruiting continues remains unknown since the size and quantity of latrines are indefinite and because the process and duration of excrement decomposition are unknown. Moreover, mushroom fruiting is affected by the annual climate.

### 2.3 Excavations and material collection

Excavations to examine the subsurface features were performed at the 91 sites and repeated at some sites for continued observations, totalling 141. Nests consisting of fallen leaves were found near mushroom fruiting points, usually at a depth of <50 cm (1 m at most). Digging was performed from the side at a distance of ca. 1 m on the same contour of the fruiting points, which should identify the ground inclination in the soil profile, and from the direction that provides the best daylight for the profile. During excavation, general nest features such as appearance, tunnel system, tunnel diameter, excrement deposition, and ‘food storage’ (Gorman and Stone, 1990) were examined. After completing digging, the soil profiles around the nest and tunnels were photographed. Materials that might be relevant were collected from any pertinent features encountered in the course of excavation, such as the nest, deserted latrines (where the soil had been colonised by mushroom hyphae and fine tree roots), and current latrines (where fresh excrement had accumulated). Furthermore, the sites’ environmental conditions, including topography and vegetation, were recorded. In rare cases, the inhabiting mole was captured, in accordance with Japanese Laws.

All materials collected by Sagara, except mushroom specimens in Europe, were deposited at the Kyoto University Museum. European mushroom specimens were deposited at the institutions of their respective countries, including the Royal Botanic Gardens Herbarium in the UK, the University of Oslo Fungarium in Norway, and Eidg. Forschungsanstalt WSL in Switzerland, or at the collectors’ personal herbaria. Mushroom specimens collected by Kasuya were deposited at the National Museum of Nature and Science, Tsukuba, Japan. Other materials such as ectomycorrhizal samples and mole’s nests excavated underneath the mushrooms were housed in Kasuya’s personal herbarium at the Department of Biology, Keio University.

### 2.4 Identification of the nest inhabitant and causal animal

Unless juveniles or young were present, the excavated nest was empty at that time since the inhabitant had either escaped from or already abandoned it. This made the identification of nest inhabitants difficult. Therefore, we studied hair characteristics to solve this problem and identify the nest inhabitants because hairs were abundant in and around the nest. General nest features (**Section 2.3**) were also compared. The tunnel diameters of Japanese moles were compared to the standards reported by Sagara (1998b).

Hairs were acquired from the collected materials. Those from the nest were considered to have fallen directly off the animal’s body by natural moulting or grooming, and those from latrines to have been swallowed during grooming. In this identification, we referred to the keys provided by Day (1966) and Teerink (1991) that apply to Japanese material (Sagara, 1986; Sagara et al., 1993b, 2008a). We found that: (1) mole hair, shrew hair, and mouse hair can be distinguished by the Day and Teerink keys; (2) within the subfamily Talpinae, the tribes Urotirichini and Talpini can be distinguished by the shape of the terminal section of awns, which is slightly wavy in *U. talpoides* (Sagara et al., 1981; for hair morphology and terms see Sagara [1986] and **Figure 4A**); (3) within the tribe Talpini, *O. mizura* can be distinguished from *Mogera* species by the scale pattern in the straight guard hair since its hair has narrower and pointed scales (Sagara, 1998b; **Figure 4B**).

**Figure 4.**
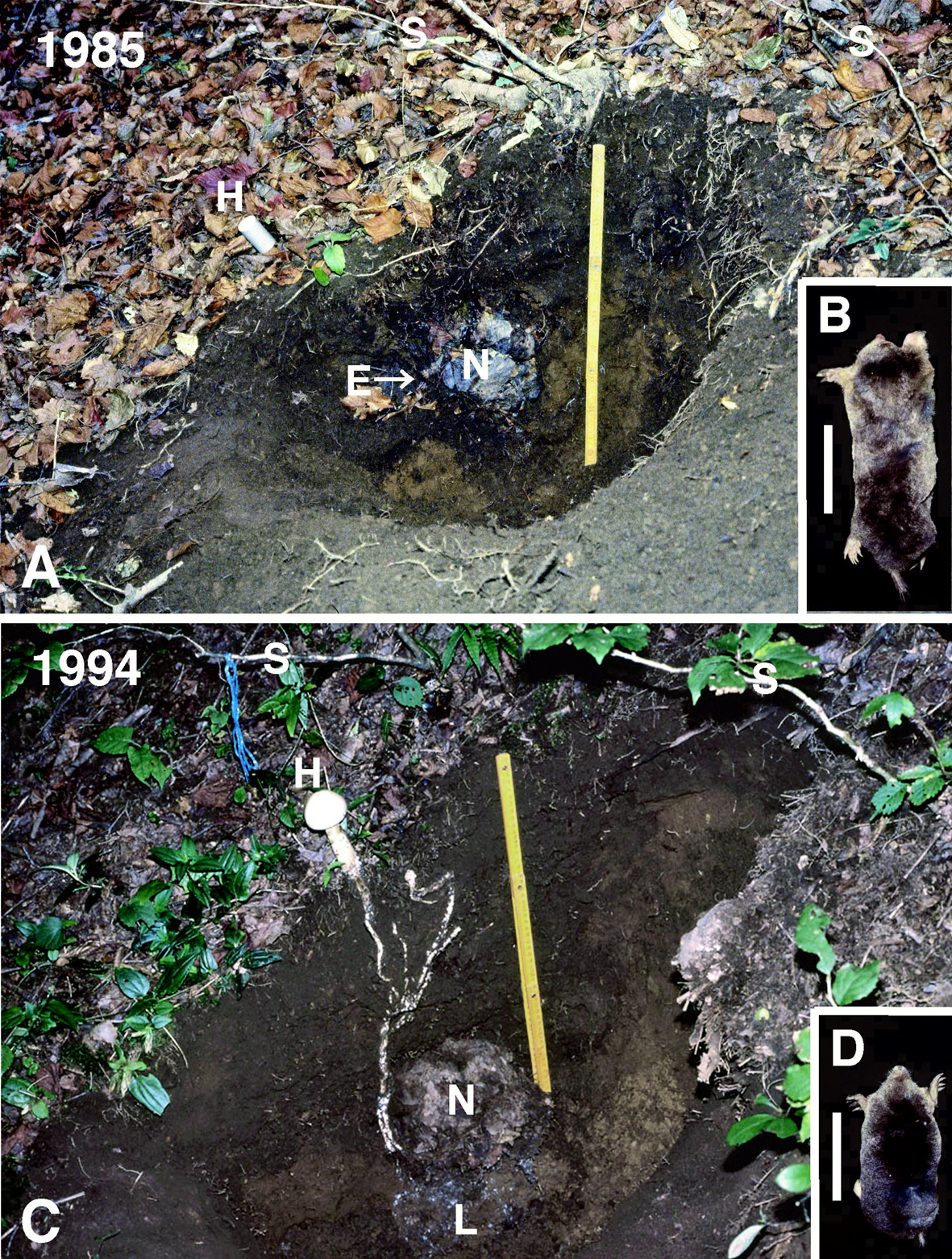
Hair types and scale pattern in Japanese moles as the material for identification of empty nests. (**A**) A group of hairs obtained from the skin of *M. wogura* shows the occurrence of three types of hairs, straight guard hairs (s), awns (a), and fur hairs (f); the bar denotes 1 mm. (**B**) Scale patterns in the straight guard hairs of three Talpini species from central Japan: **a**, *O. mizura*; **b**, *M. imaizumii;* **c**, *M. wogura*; **d**, hair from an empty nest, showing the same scale pattern as **a** and identifying the nest inhabitant as *O. mizura*; the bar denotes 50 µm. **A** from Sagara (1986), **B** from Sagara (1998b); reproduced with permission.

This identification remained unfinished for some Japanese Talpini species due to a lack of evidence and was considered provisional in the data presentation (**Table 1**, footnote). In the European examples we studied, once it was determined that the animal species in question was a mole, no further identification work was required since only one mole species, *T. europaea*, is present in that region of Europe.

Mole and wood mouse (*Apodemus*) nests appear similar in terms of material and structure, but there are approaches to distinguish them other than hair. The tunnel system is less developed in wood mouse nests, with no tunnel beneath the nest chamber (Sagara et al., 1988, 1993b, 2006). The wood mouse nests in Europe (**Table 1**) were accompanied by remains of stored food, such as beech seeds, but the mole nests, both in Europe and Japan, were not. The identification methods for shrew nests and badger setts are omitted.

The causal animal responsible for the occurrence of mushrooms by depositing excrement (urine and faeces) was determined with caution since it might differ from the identified nest inhabitant. There is an example where the nest occupant appeared to change from a mole to a mouse (or mice; Sagara, 1995a), but we exclude this here because it was unconfirmed. There are a few examples of the nest inhabitant changing from one species to another within the Japanese Talpini (**Section 3.7**). The hair and tunnel characteristics were again considered essential for identification in such circumstances.

In one example from England and another from Sweden, vole hair and mouse hair, respectively, were abundant in and beside the excavated mole nests, but they were considered to have come from bodies devoured by moles (Sagara, 1989; Sagara and Marstad, unpublished).

## 3. New findings on mole natural history

### 3.1 Overview of the mole-mushroom relationship

An example of the soil profile indicating a forest-dwelling mole nest and *H. sagarae* fruit bodies growing out of nearby abandoned mole latrines is shown in **Figure 5A**. Ectomycorrhiza formation, a symbiosis between mushroom hyphae and tree roots, at such latrines is shown in **Figure 5B-C**. A diagram of this tripartite biological relationship between moles, mushrooms, and trees in which mole excrement is transformed and translocated is presented in **Figure 6** (**Section 3.3**). These examples also hold for *H. radicosum* in Europe.

**Figure 5.**
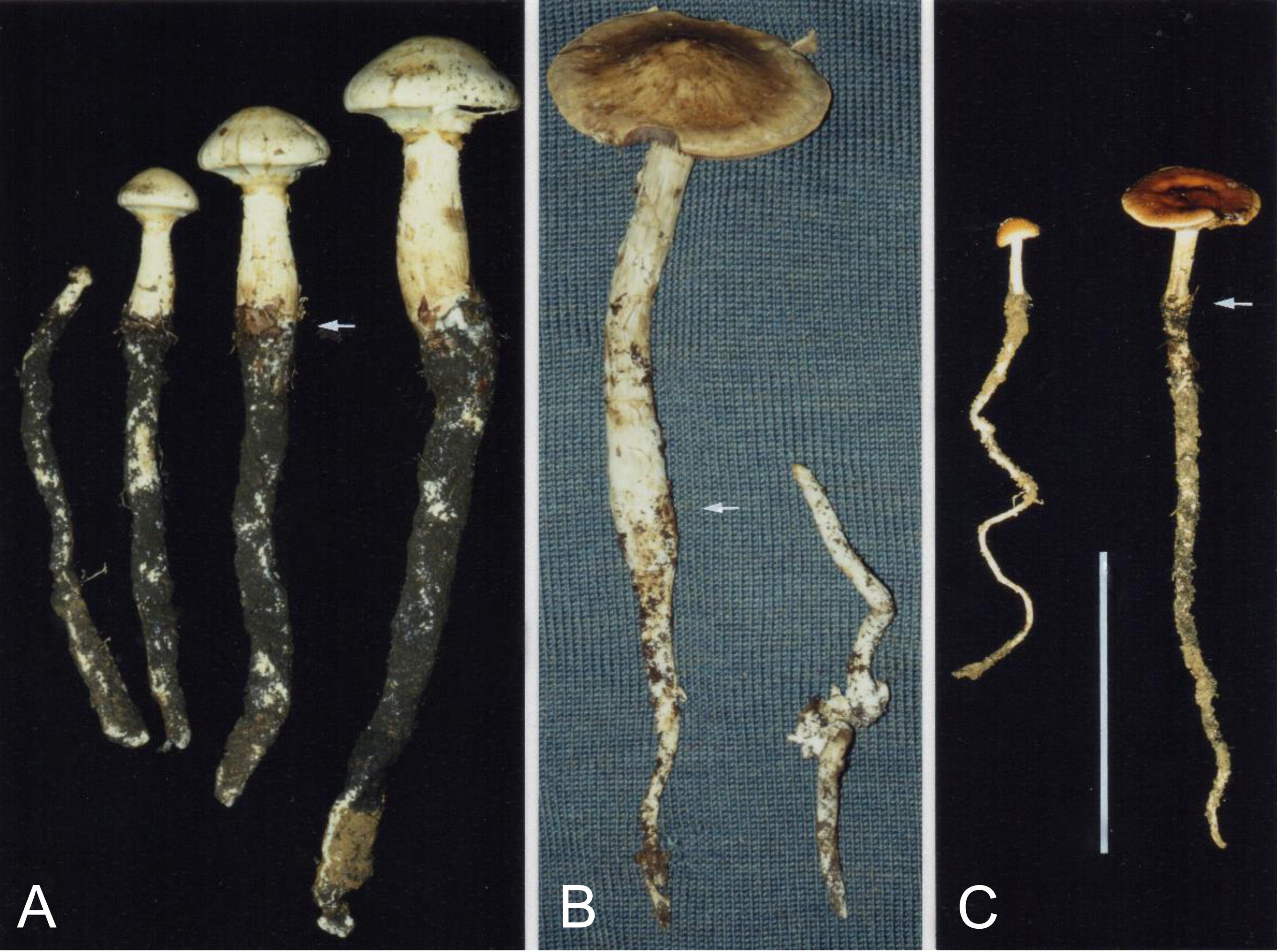
Observed features of the relationships among moles, mushrooms, and trees that have arisen from *O. mizura* mole excretion. (**A**) Soil profile showing the mushroom *H. sagarae* fruiting (H) on deserted mole latrines (L). Key: S, rooting (subterranean) stipes of the fruit bodies; N, nest; the bar denotes 10 cm. (**B**) Masses of mushroom hyphae (appearing white) and fine tree roots colonising the soil of a deserted mole latrine; the bar denotes 10 mm. (**C**) Photomicrograph of a transverse section of the fine roots that have formed symbiotic ectomycorrhizas, as characterised by the presence of a fungal sheath (F) and Hartig net (H). Key: E, epidermal root cell; Co, cortical cell; Cc, central cylinder; the bar denotes 10 µm. Original photographs: **A** from Kutsuki, Shiga Prefecture, Japan, 260 m alt., 9 October 2001; **B** from Ashiu, Kyoto Prefecture, Japan, 670 m alt., 9 October 1987; **C** root material from Ashiu, 735 m alt., 18 November 1990 (see **Fig. 14** in Sagara [1995] for further information).

**Figure 6.**
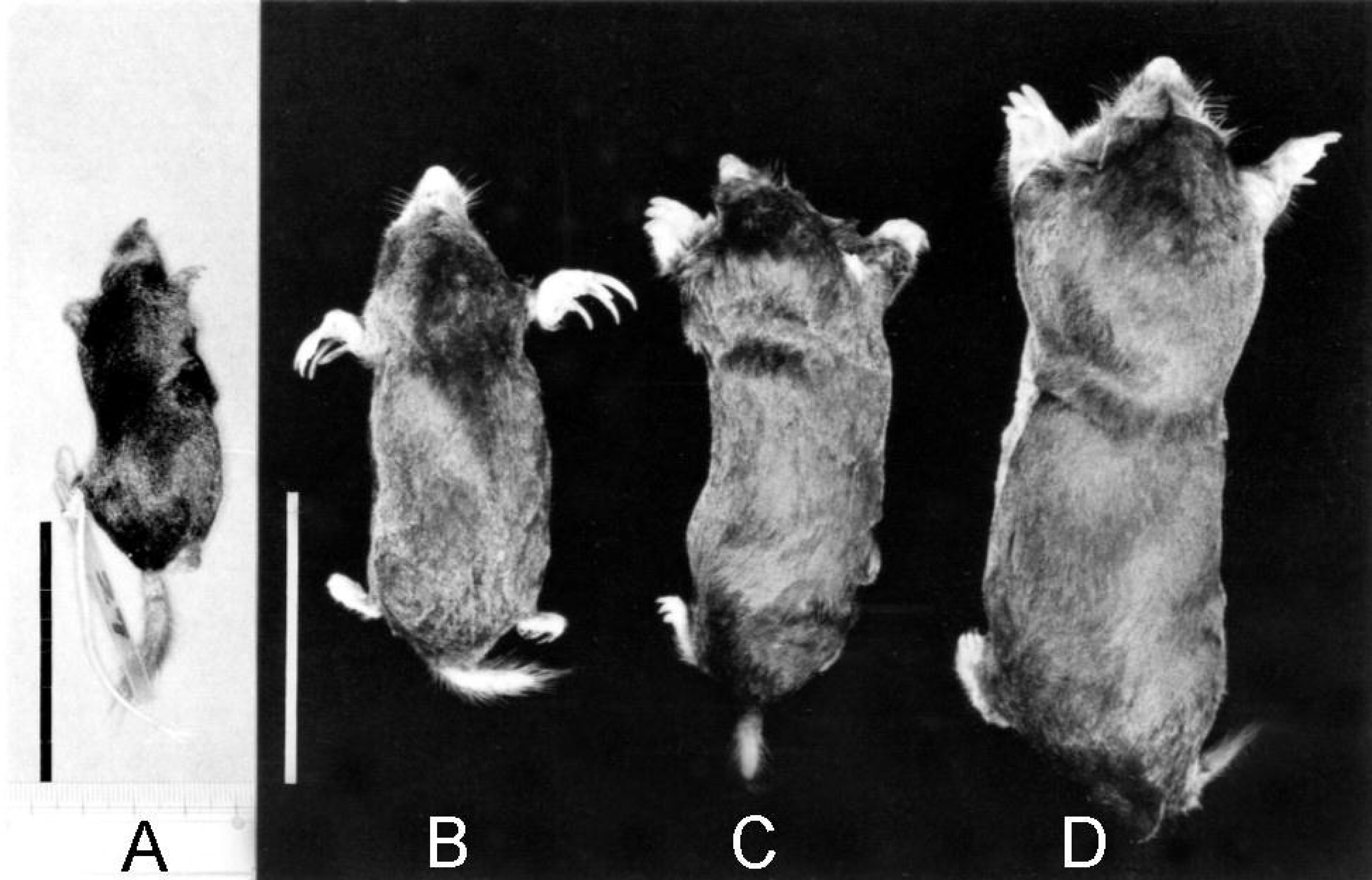
Diagram of the observed relationship among moles, mushrooms, and trees, hypothesised as a habitat-cleaning symbiosis, in which mole excrement is cleaned away by mushroom hyphae and fine tree roots forming mycorrhizas (ectomycorrhizas). Here, moles rely on mushrooms and trees to continue living in the same nests. Modified from Sagara (1999). For further explanation and discussion, see **Section 3.3**.

The mole in this relationship was, in rare cases, replaced by other mammals such as the wood mouse *Apodemus* (**Table 1**). However, in Honshu, the main island of the Japanese Archipelago where *Apodemus* species ecologically equivalent to the European species exist, all 77 examples of *H. sagarae* and *H. danicum* occurrence studied were associated with moles; there are no examples associated with mice. Therefore, most cases of these *Hebeloma* species were due to moles, and research results have concentrated on moles. Talpidae and Soricidae are also found in North America, but this relationship has not been explored there.

Tree families possibly involved in this relationship as hosts of the mycorrhizal symbiosis include Fagaceae, Betulaceae, and Salicaceae, based on field and microscopic observations (Sagara, 1995b, 1999; **Table 2**). These families are known to be ectomycorrhizal (Smith and Read, 1997). Pinaceae are also ectomycorrhizal, but no example of this relationship has been observed in any pure conifer forest. Intriguingly, in coniferous *Picea-Abies* forests in Switzerland, the *H. radicosum* sites studied were all confined to the rare, spotted growth of beech *Fagus sylvatica* trees (**Figure 3A**), suggesting a strong association between this mushroom species and ectomycorrhizal broad-leaved trees. In addition to mycorrhizal symbiosis (**Section 3.3**), the supply of broad leaves as nest material may be an essential aspect of mole life (**Section 4C**).

The occurrence of *H. danicum* on mole latrines was relatively rare. This species, being an ammonia fungus (Sagara, 1975), occurs not only on mole latrines but also on various animal corpses and excrement after their decomposition, even in coniferous forests (Sagara, 1975, 1992, 1995b; Sagara et al., 2008c). Its occurrences on mole latrines, including possible incidents in coniferous forests, may have often been overlooked since this mushroom is inconspicuous. *Hebeloma radicosoides* Sagara, Hongo, & Y. Murak., another ammonia fungus that occurs on various animal wastes, including mouse excrement (Sagara et al., 1993b), is highly similar to *H. sagarae* in appearance but has never been found on mole latrines (Sagara et al., 2000, 2008c).

Recognition of the tripartite association for forests does not preclude the possibility of moles nesting in open land such as arable fields, where cleaner organisms other than particular mushrooms and trees could operate. However, there has neither been reported nor observed such long-term nesting described in **Section 3.8** on open land.

In addition, Watling (1978) observed in England that *H. radicosum* was ‘intimately connected to the urine-soaked fur and soil of the tunnel of a small mammal’, but he did not refer to the mammal’s species nor its nest. His observation was published shortly before that of Sagara (1978), but as he recognises, had been prompted by Sagara.

### 3.2 Latrine making by moles

‘Latrine’ is used here to refer to the part of the ground, with or without any tunnel enlargements or depressions, in which excrement (urine and faeces) has been repeatedly deposited and accumulated in large amounts. The word ‘midden’ has also been used in some of our articles to refer to latrines. We have used many examples in our previous articles to explain the cause of the *Hebeloma* occurrence. A thorough feature of the latrine making is shown in **Figure 7**, and further examples can be found in **Figures 5A, 14, 16B+D**, and **17C**.

**Figure 7.**
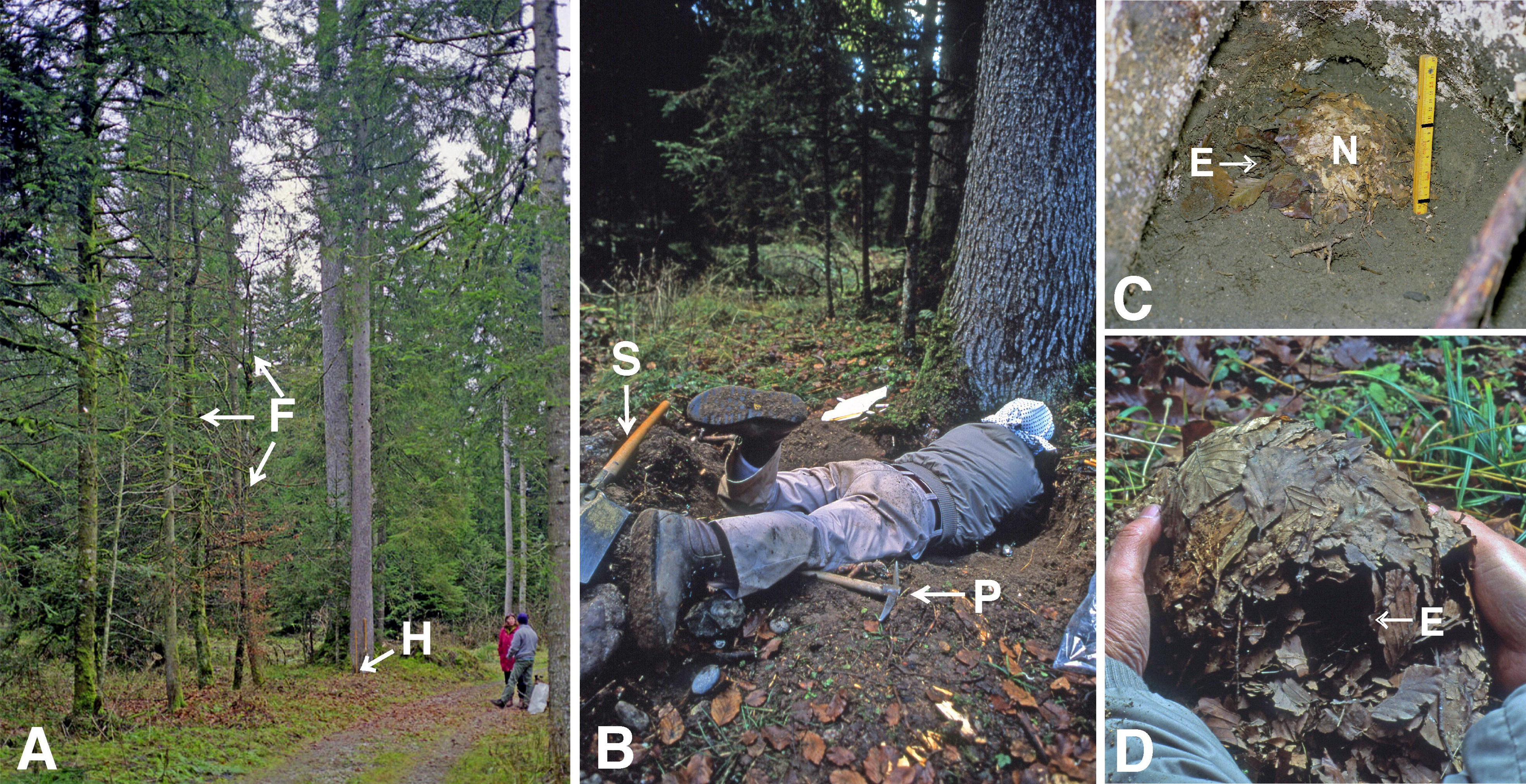
*Oreoscaptor mizura* latrines. (**A**) A new latrine (L) found in front of the new (reconstructed) nest (N). Key: E, entrance to the nest; T, a tunnel leading to the latrine. (**B**) Excrement deposit (D, dark-coloured) removed from the latrine in A. Thick lines on the folding scale in A and B are at 10-cm intervals. (**C**) Soil block containing a newer latrine (NL) formed on an older latrine (OL), with the latter having been colonised by fungal hyphae and fine tree roots forming ectomycorrhizas; the bar denotes 10 cm. For further information, see **Figs. 8, 9, and 14** in Sagara (1999). Photographs are original.

Talpini mole latrines are usually located within 50 cm and rarely as much as 1 m away from the nest. However, *U. talpoides* latrines are often scattered and can be found even 3 m away from the nest (**Figure 8**). Moles appear to relocate their latrines little by little in the vicinity of their own nests, as indicated by the change of mushroom fruiting points (**Fig. 11** in Sagara, 1999; cf. **Figure 15**).

**Figure 8.**
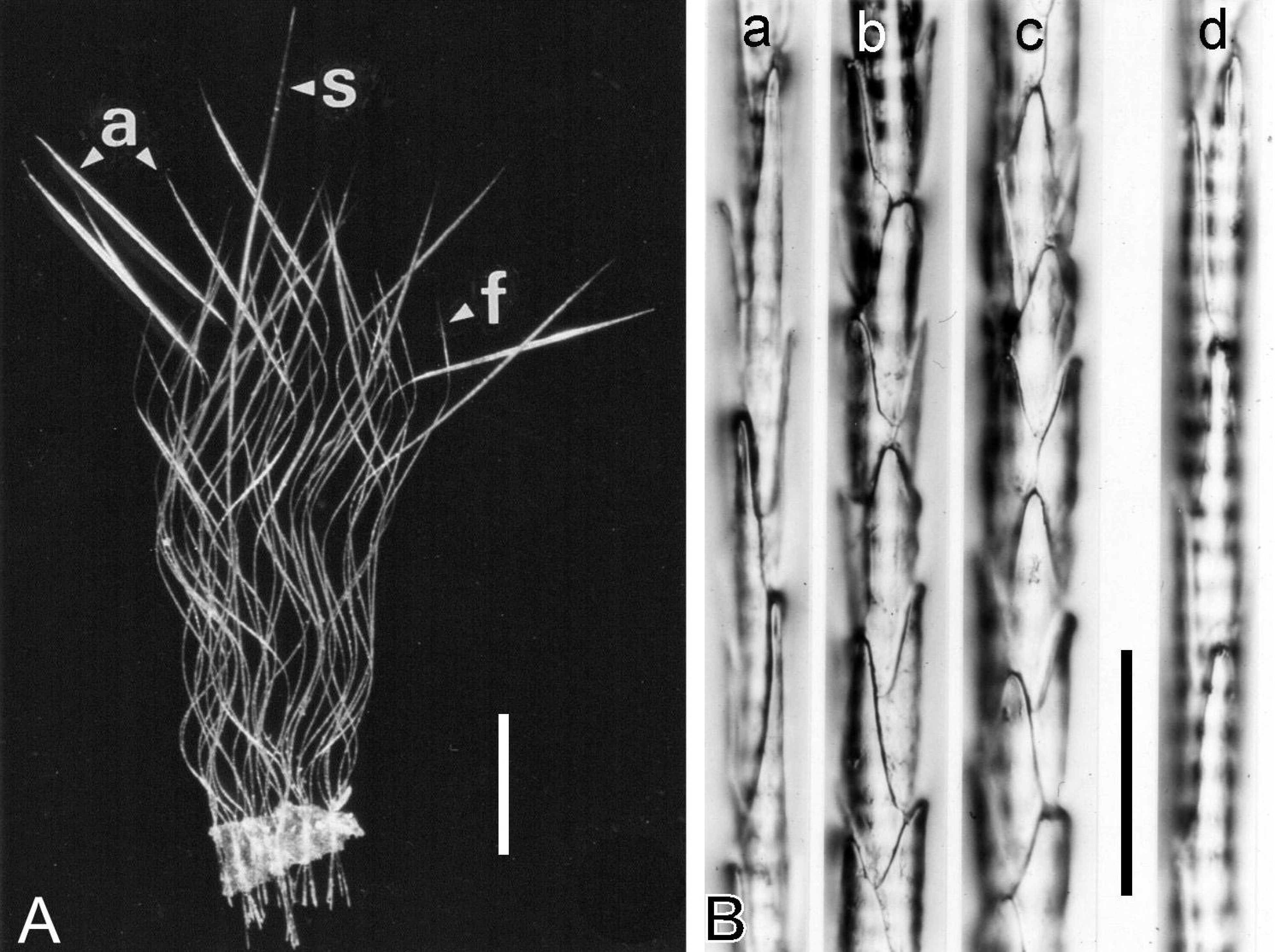
Map showing an example of the scattered latrines of *U. talpoides* as indicated by the scattered occurrence of *H. sagarae* and *H. danicum* fruit bodies. Ground inclination 35–44° NNW. Key: F, a footpath in the forest; N, nest; Hr, *H. sagarae* (previously *H. radicosum*); Hs, *H. danicum* (previously *H. spoliatum*): dots, positions of the fruit bodies. Modified from Sagara et al. (1981).

**Figure 9.**
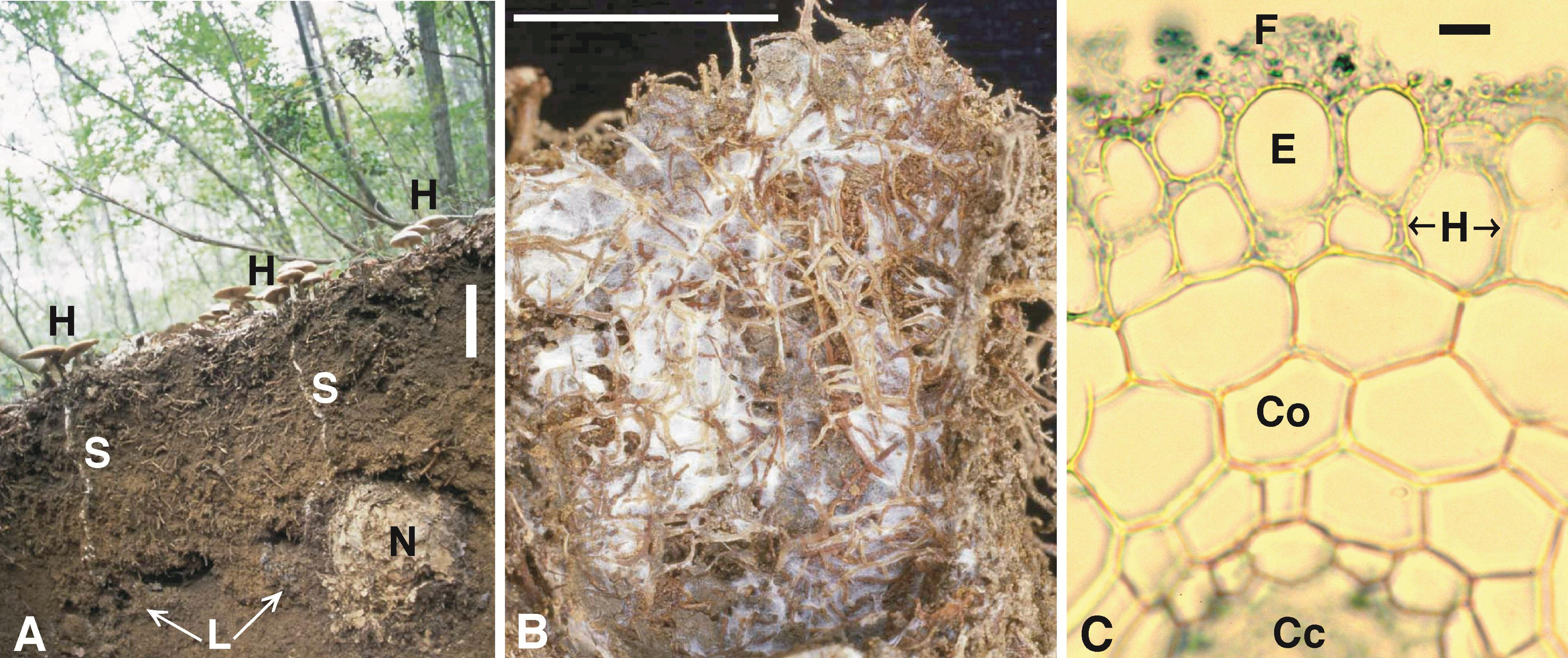
Examples of newly reported *U. talpoides* and *O. mizura* nests in this study. (**A**) Soil profile containing a *U. talpoides* nest under the fruiting points of *H. sagarae* (sticks H). Key: G, ground surface; E, entrance to the resting cavity; T, tunnel; the folded scale is 20 cm long. (**B**) Soil profile containing an *O*. *mizura* nest that was newly constructed after removal of the previous nest at the same location 24 days earlier. Key: G, ground surface. The entrance to the resting cavity was on the opposite side and cannot be seen in this photograph. (**C**) A long-used nest of *O. mizura* with a dense root cover (nest placed on a polyethylene bag; see text for root cover). Key: E, entrance to the resting cavity. The bars denote 10 cm. From Sagara (1999), reproduced with permission.

**Figure 10.**
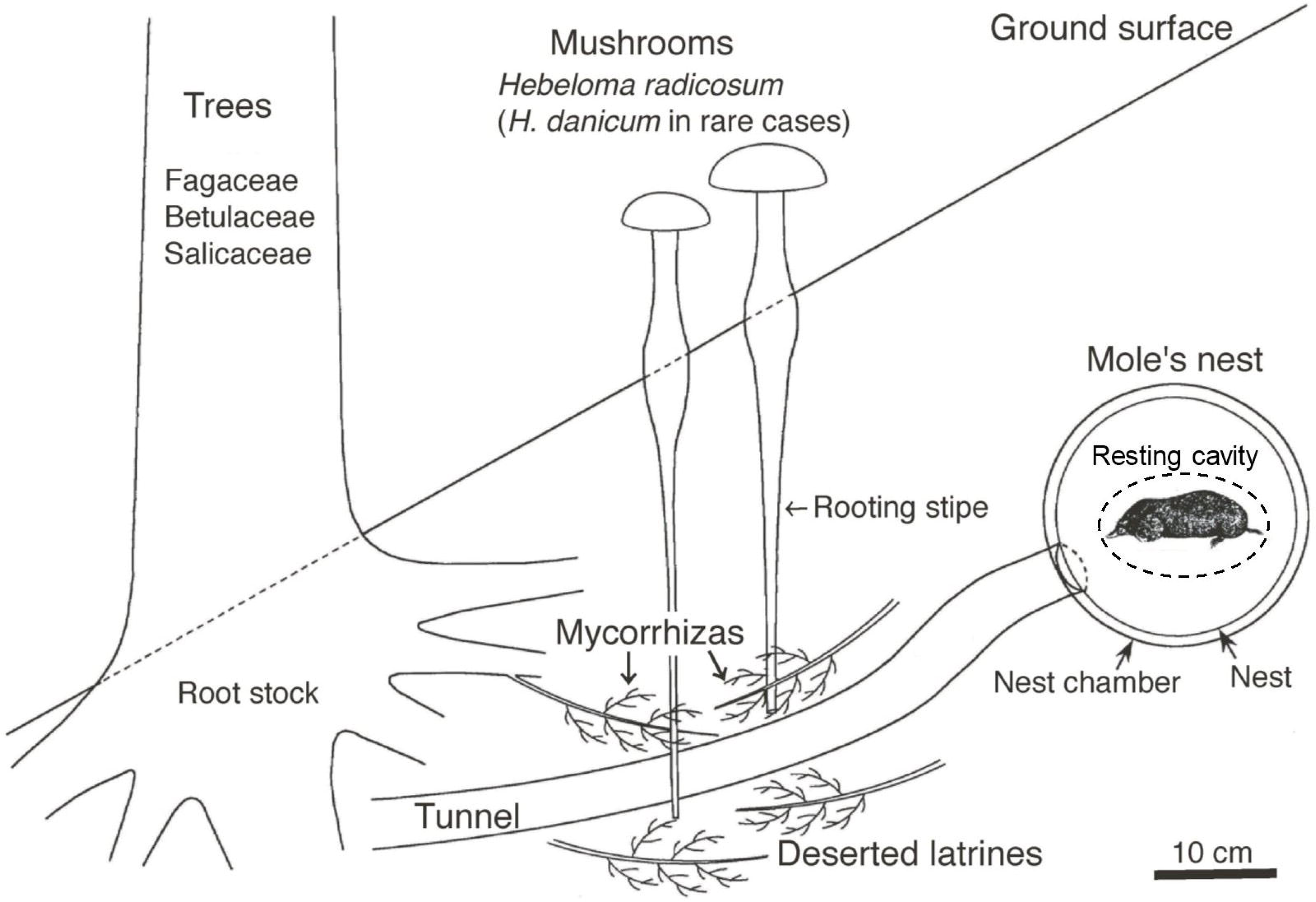
The internal structure of *M. imaizumii* nests shows the presence of a single entrance (E) and a closed resting cavity (C). (**A**) X-ray computed tomography (CT; vertical scan) of the central part of a nest. (**B**) Outline of a nest, obtained by sectioning it vertically at its centre. Key: W, wall composed of fallen leaves. The bars denote 5 cm. From Sagara (1998b), reproduced with permission.

**Figure 11.**
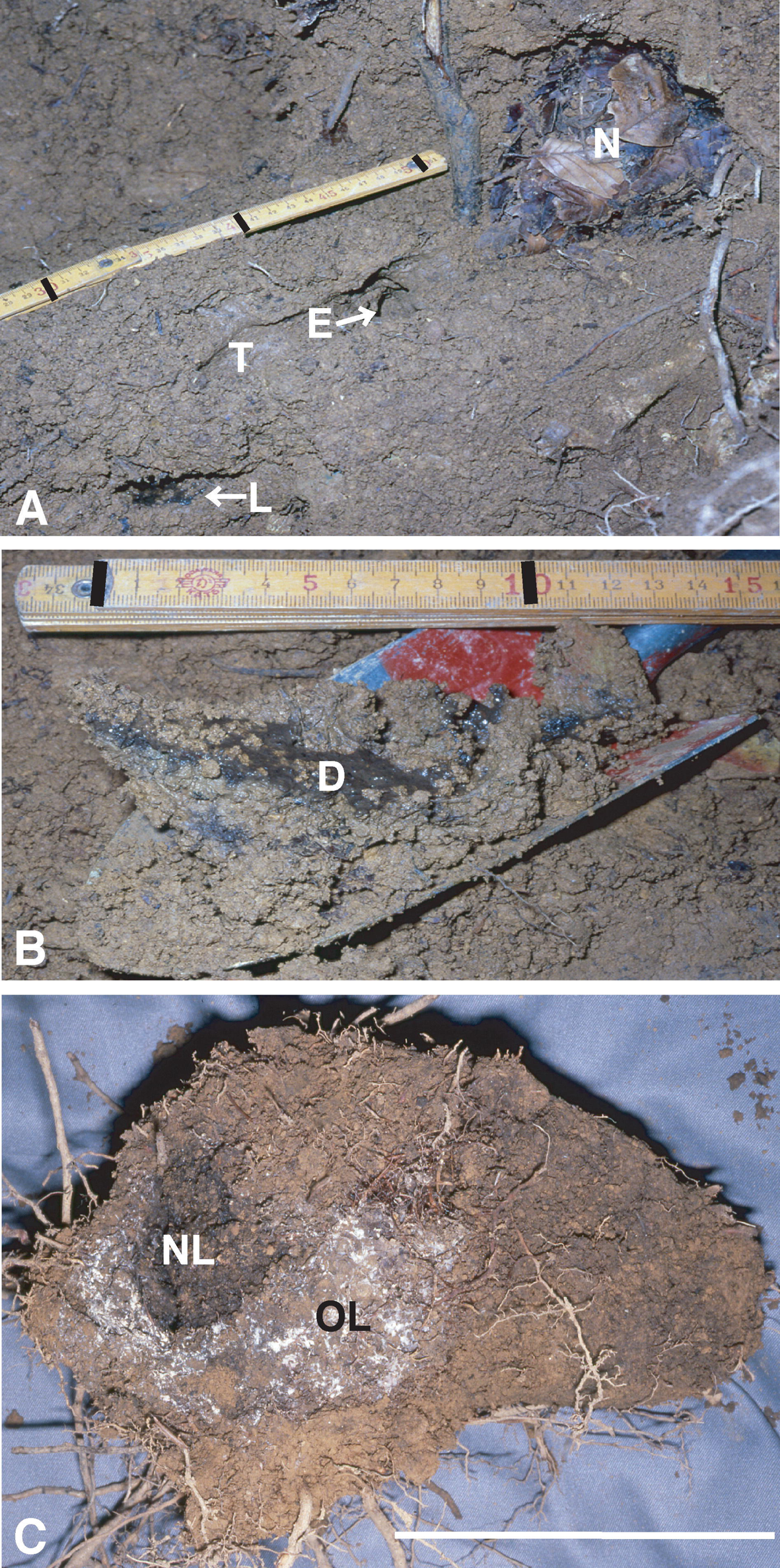
Distribution of mole nesting sites within a forest stand, as indicated by the occurrence of the mushroom *H. sagarae* during one autumn season in 2000. Key: 1 or 3, *O. mizura* nesting sites; 2, *M. wogura* nesting site; P, path. Surface distances between the nesting sites were ca. 41 m between sites 1 and 2, ca. 44 m between sites 2 and 3, and ca. 17 m between sites 3 and 1. Locality: Kogaino, Yamazaki-cho, Shiso-shi, Hyogo Prefecture, Japan, 660 m alt. Photograph: 16 December 2000. Note: This photographing was possible by the forest management particular to this area, where the undergrowth had been clear-cut, and by information from a local amateur mycologist who had been intensely watching mushrooms there. Sagara, unpublished.

**Figure 12.**
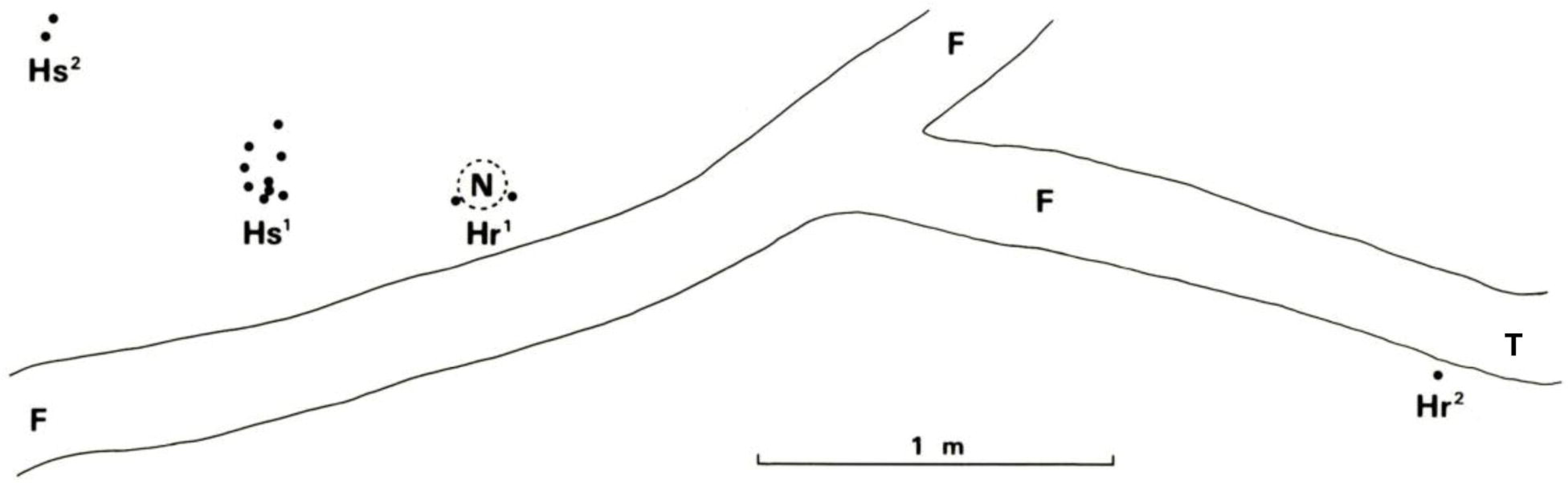
Nesting by different Talpini species in close proximity to one another in mountain habitats, as indicated by the coincident occurrence of *H. sagarae* (and *H. danicum* in **B**) at two sites marked in the maps. Contour interval: 10 m in **A** and **B**; 5 m in **C**. *E. mizura* in **A** and **B** now represents *O. mizura*. From Sagara (1999), reproduced with permission; see this article for further details.

**Figure 15.**
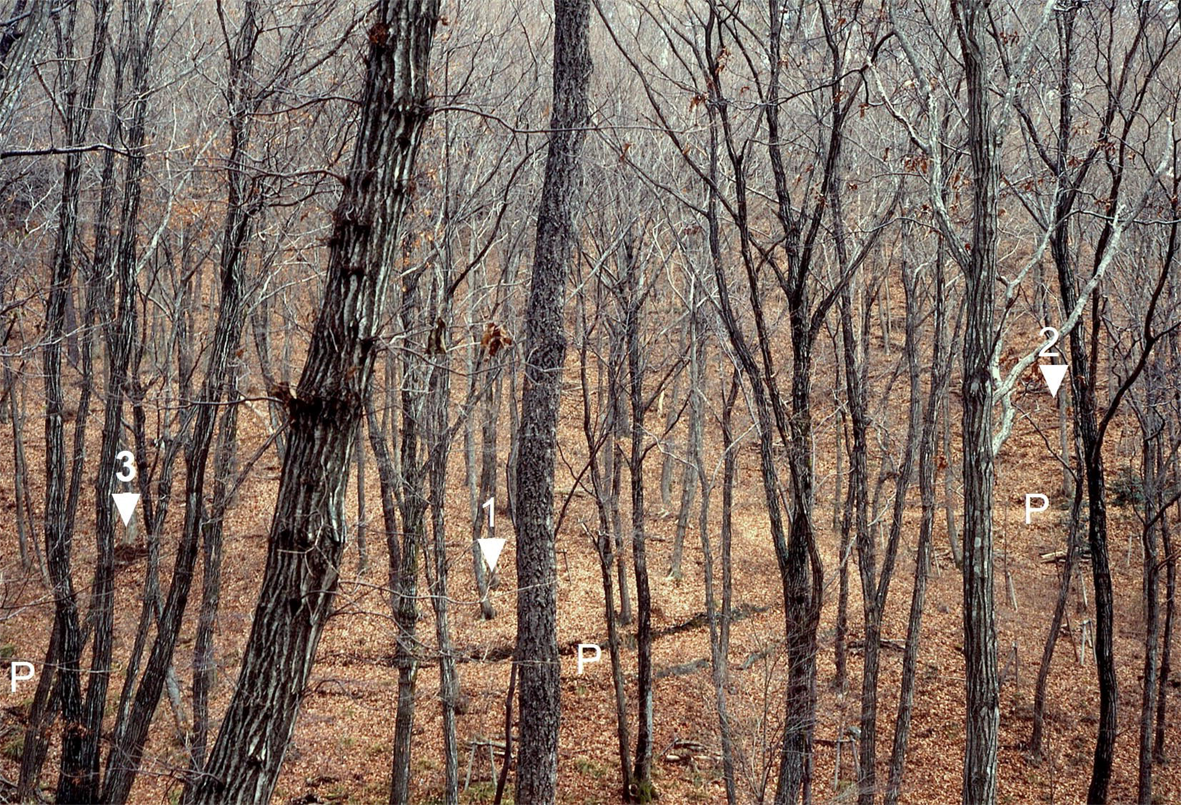
Site of long-term nesting by *O. mizura* as documented in **Table 3**, with marks of the mushroom *H. sagarae* fruiting points and of nest sites that changed between 1981 and 1999 under repeated excavations and nest removal. The mushroom fruiting at this site was first discovered in 1977, but excavation was not conducted during 1977−1980. Marks 81−98 indicate fruiting points in 1981−1998, respectively. Marks N81−N84 indicate underground nest sites found by excavation in 1981−1984, respectively. Mark N85/ N89 indicates the nest site used from 1985 to 1989, as revealed by excavations in 1985 and 1989. Mark N99 indicates the nest site used from 1989 to 1999 found by excavation in 1999. Key: F, fallen tree. The folding scale is 1 m long, with divisions at 10-cm intervals. Photograph 8 July 1999. Modified from Sagara (1999).

**Table 3.**
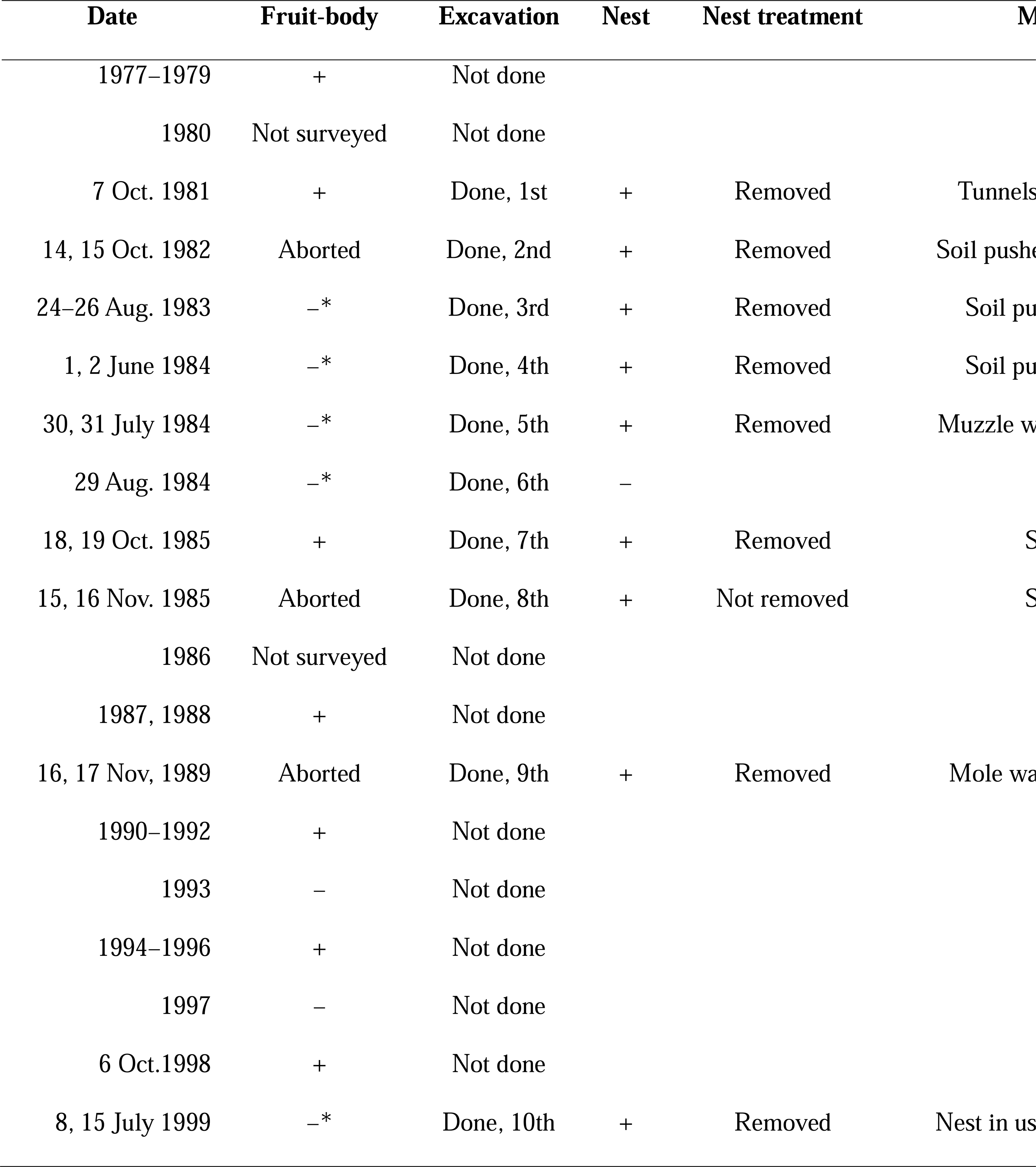
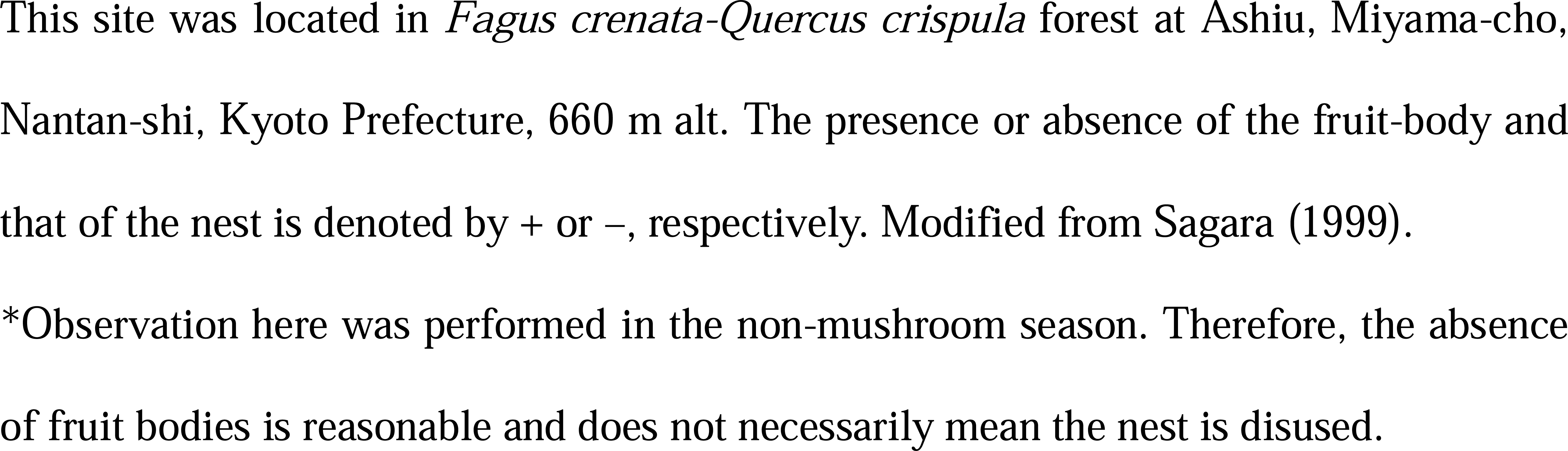
Long-term observations on an *Oreoscaptor mizura* nesting site with repeated *Hebeloma sagarae* occurrences (see **Figure 15**)

Mole urine is known to contain scents secreted from preputial glands beneath the skin. Scent marking with urine for olfactory communication in moles has been documented (Gorman and Stone, 1990), but latrine making has not been discussed in this context. The latrine contains urine and dung, which yields a particular odour from themselves and their decomposition. We have not explored the olfactory effects of the mole latrine but have had the impression that some observations in our study may be relevant. For example, the scattered latrines and faece-trodden tunnels in *U. talpoides*, the latter being quite extensive around the nest (Sagara et al., 1981; **Figure 8**), may also act as scent marks. Indeed, the latrines under the *H. sagarae* fruit bodies in **Figure 8** (Hr^2^; cf. Fig. 24 in Sagara et al., 1981) strongly suggest the significance of home range marking because they were located at the turning point of the tunnel that led ca. 3 m away from the nest. In addition, the return home of Talpini moles from foraging could be guided by the olfactory effects of the latrines beside their nests.

### 3.3 Mycorrhiza formation in latrines: habitat-cleaning symbiosis

See **Figure 6** for a general idea discussed in this section. When a mole latrine becomes old or deserted, mushroom hyphae and fine tree roots simultaneously colonise it and form ectomycorrhizas (Sagara, 1995b, 1999; Mikami, 2015; Maeno et al., 2016; Kasuya et al., 2017; **Figures 5B-C and 7C**), a type of mycorrhiza that forms a symbiotic relationship between mushrooms and plants (Smith and Read, 1997). *Hebeloma radicosum* and *H. spoliatom* (probably *H. danicum*) are known to form an ectomycorrhizal association with birch *Betula* spp. in axenic culture (Giltrap, 1982). The identity between *H. sagarae* fruit bodies appearing aboveground and the hyphae forming ectomycorrhizas in the mole latrine underground has been established by molecular analyses (Maeno, 2000; Mikami, 2015; Maeno et al., 2016; Kasuya et al., 2017). Kasuya et al. (2017) further identified the plant symbiont as *Quercus serrata* (Fagaceae) for the ectomycorrhizas they collected from a mole latrine.

Excrement decomposition and turnover in the mole latrine appear to proceed as follows. Excrement is first decomposed mostly by bacteria and small invertebrates, although certain ammonia fungi such as *Amblyosporium botrytis* Fresen. may also be involved (Sagara, unpublished). Next, the mushroom hyphae and tree roots, forming ectomycorrhizas, absorb and utilise the decomposition and turnover products. Then, the absorbed substances are transformed in the hyphae and tree roots, and translocated above-ground, partly by mushroom fruiting and partly by tree growth. Consequently, the mushrooms and trees act as the cleaners of mole latrines. Therefore, moles benefit from those mushrooms and trees, despite not relying on them for food (Sagara, 1995b, 1999).

This relationship differs from the ‘cleaning symbiosis’ proposed by Eibl-Eibesfeldt (1970) since the mushrooms and tree symbionts clean the animal’s habitat, not its body. Therefore, this relationship has been proposed as a ‘habitat-cleaning symbiosis’ (Sagara, 1999; Sagara et al., 2008c). The mutualism here may further be explained as follows (Sagara, 1995b): The plants provide habitat, niches and, by forming mycorrhizas, waste cleaning for the animals, and energy and probably other vital compounds to the mushrooms (fungi). The animals provide nutrients to the plants and the fungi, ploughing and aerating the soil for them. The fungi provide nutrients and growth regulators to the plants and waste cleaning for the animals. Indeed, waste cleaning would be essential for enabling moles to continue nesting for an extended period (**Section 3.7**).

The mutualism hypothesised here would apply to cases where excrement or dead bodies of other animals, including humans, are naturally decomposed in forest habitats (Sagara, 1995b, 2000; Sagara et al., 2008c).

### 3.4 Undescribed nests unveiled, and the nest structure clarified

Mole nests, especially those of Japanese species, have scarcely been observed. The nests of *U. talpoides* and *O. mizura* were first documented by Sagara et al. (1981, 1989) (**Figure 9**). Information on the nests of *M. imaizumii*, *M. wogura*, and *T. europaea* was increased (Sagara, 1978, 1980, 1998b, 1999; Sagara and Abe, 1993).

Moles are known to construct their nests using fallen leaves and dry grasses, but the structure of the nests has not been correctly described. They have often been described as having two or more entrances, as illustrated in **Figure 2.3** in Gorman and Stone (1990). Sagara (1998b) observed (**Figures 6** and **10**) a uniform structure that has a closed resting cavity at its centre (‘nest cavity’ in Sagara [1999]) and a single entrance leading to it. This structure applies to all studied talpid species, including *T. europaea* (Sagara, 1989, 1998b; Sagara and Abe, 1993; Sagara et al., 1989; **Figures 3C-D, 9A+C, 14, and 17A**). The only relevant description of talpid nests was provided by Eadie (1939) on *Parascalops breweri* (Bachman) nests in North America, where he observed an ‘inner cavity’ in the spherical nest and ‘a loose passageway through the otherwise compact wall on the side’.

The long-used nest is usually enveloped with a layer of fine tree roots that have likely developed to absorb the nutrients from decomposing nest material (**Figure 9C**).

The nest chamber is a subterranean room in which a nest is situated (**Figure 6**) and is poorly understood. We found that the nest chamber also has only one entrance that is directly connected to the resting cavity of the nest (Figs. 22 and 23 in Sagara [1999]; **Figure 6**). The nest chamber of *T. europaea* has been described as having two or more entrances (**Figure 8** in Godfrey and Crowcroft [1960]; **Figure 2.3** in Gorman and Stone [1990]). Similar descriptions also prevail in the literature on North American moles. Indeed, it often appears that multiple tunnels lead away from the nest chamber. However, these tunnels, except the one leading to the resting cavity, are dead ends. We once incorrectly described the nest chamber as having multiple entrances (**Fig. 6** in Sagara [1978]; Sagara et al., 1981).

At several *U. talpoides* sites, the nest itself could not be found, but the species’ involvement could easily be confirmed by hair and tunnel characteristics (Sagara et al., 1981; **Section 2.4**). One possible explanation was the positional relationship between *U. talpoides* latrines and the nest. It would be difficult to locate the nest if the latrines are scattered (**Figure 8**), and if mushrooms occur only on latrines distant from but not near the nest. Another possibility is that *U. talpoides* may nest in unknown ways, supposedly, by not making such underground nests as shown in **Figure 9A**.

### 3.5 Mole distribution revealed, not by capture, but by nest identification

Identifying nest inhabitants without capturing, as in this study, has unveiled new locations in the distributions of some mole species (Sagara, 1999: Nakai et al. 2016). We found *O. mizura* in previously unreported areas of Okayama, Hyogo, Kyoto, and Shiga Prefectures (Sagara et al., 1989, 2008b; Sagara, 1999), showing that it is distributed more widely than previously thought, in the mountains and hills of western Honshu. In addition, we found *M. imaizumii* nests in Shimane, Hyogo, Kyoto, Shiga, and Aichi Prefectures (Sagara et al., 1993a; Sagara, 1998b, 1999, unpublished data), where the presence of this species is not well recognised.

*Oreoscaptor mizura* has been considered a highland animal, as reflected by its common name, Japanese mountain mole. However, we found its nesting sites at lower altitudes and, at one (260 m above sea level; **Figure 5A**), captured the inhabitant (Sagara et al., 2008b). While there have been reports of dead *O. mizura* at lower altitudes, our study provided direct evidence of this species living there.

### 3.6 Presence of different Talpini species in the same area

Japanese Talpini species, particularly *M. imaizumii* and *M. wogura*, are often considered allopatric, based primarily on data from agricultural land on alluvial plains (Imaizumi and Imaizumi, 1970; Abe, 2010, and papers cited therein). Conversely, we have often observed that *O. mizura*, *M. imaizumii*, and *M. wogura* live close by within the same stand in montane forests (Sagara et al., 1989; Sagara, 1999; Sagara and Fukasawa, 2014; **Figures 11, 12, and 17**). ‘Mixed distribution’ or ‘sympatric occurrence’ of different Talpini species has also recently been reported on open land, including paddy fields (Moribe and Yokohata, 2011; Sato et al., 2021). It is likely that their occurrences are not always allopatric.

An example of the distance between neighbouring nests used by the same mole species is shown in **Figure 11**. Here, two *O. mizura* nests were located approximately 17 m apart and appeared based on their aged appearance to have been used for many years during overlapping periods. However, it is unknown whether these nests were used by the same individual or different individuals. Two similar examples were reported previously, one for *O. mizura* and another for *M. imaizumii* (Sagara, 1999), while there had been no relevant reports on Japanese moles.

### 3.7 Nestlings encountered: information on breeding

#### Oreoscaptor mizura

The biology of this species is least known (Ohdachi et al., 2009). We identified its young *in situ* for the first time and brought an individual successfully into captivity (Sagara, 2009; **Figure 13**). This result provided the first information on the breeding of *O. mizura* and the first report of captive rearing a young Talpini mole that had not yet left its maternal nest and had probably not yet weaned.

**Figure 13.**
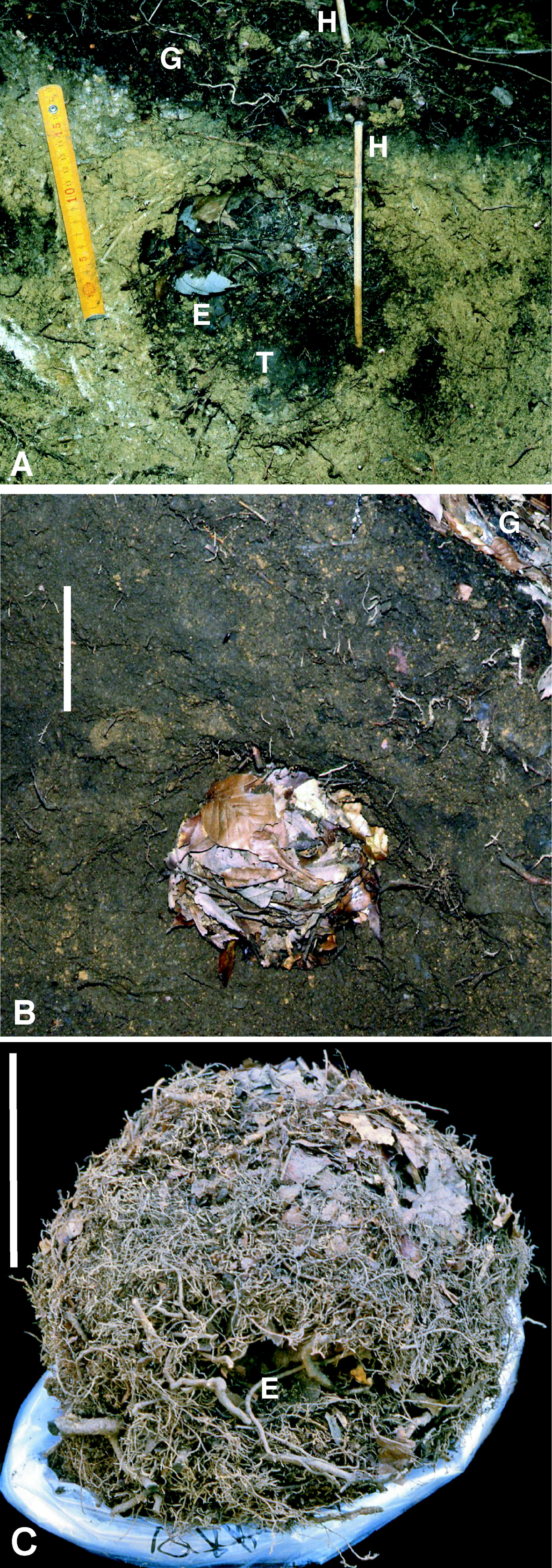
*Oreoscaptor mizura* young found *in situ* and brought into captive rearing. (**A**) Three nestlings were found after the removal of the nest ceiling. (**B**) After 19 days, the nestlings appeared close to natal dispersal; one of the two was taken into captivity at this stage. (**C**) The young mole (arrow) successfully fed a half-cut mealworm despite the mole’s teeth appearing underdeveloped. (**D**) The young mole on a human hand, appearing healthy and normal with its forefoots still delicate and likely weak; E, eye. From Sagara (2009), reproduced with permission. See also Site 1 in Sagara and Fukasawa (2014).

#### Mogera imaizumii

Nestlings of this species at similar developmental stages as with *O. mizura* were first recorded on video by Masahiro Iijima (Asia Nature Vision) and Yasuhiro Adachi (NHK, Japan Broadcasting Corporation) using a fiberscope camera at a *H. sagarae* site found by us at Kutsuki, Takashima-shi, Shiga Prefecture (Iijima and Tsuchiya, 2015). This footage was broadcast by NHK on 16 July 2006 and appears to be the first successful video record of talpid nestlings worldwide.

#### Mogera wogura

Talpini moles breed, in principle, once a year in spring (Godfrey and Crowcroft, 1960; Abe, 1968; Gorman and Stone, 1990), and *M. wogura* moles reportedly breed between April and June (Abe, 1968). An excavation following the fruiting of *H. sagarae* in October revealed the presence of *M. wogura* juveniles nearing natal dispersal (Sagara and Abe, 1993; **Figure 14**). This brood was not thought to be the second in that year but instead delayed due to the unusually cold weather that spring. Nevertheless, this result formed the first Japanese report of Talpini juveniles belonging to the same litter and approaching natal dispersal collected in autumn at their nesting site.

**Figure 14.**
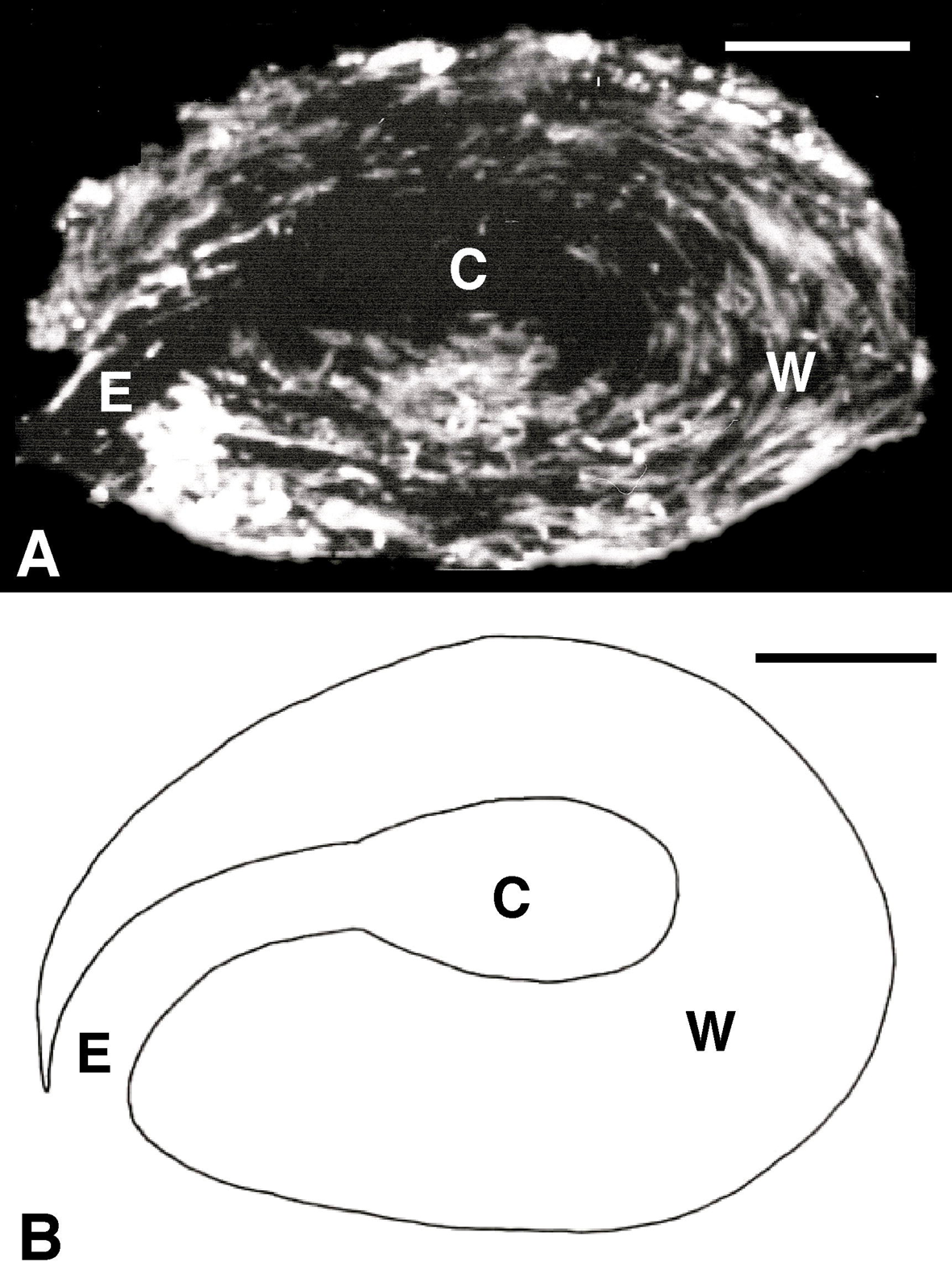
*Mogera wogura* young were found by the occurrence of mushroom *H. sagarae* in early autumn (9 October 1984), well outside its normal breeding season. The young (M) were found in situ after anesthetisation through the tunnel (T) under the fruit bodies (H), while their mother escaped from the site and anesthetisation. Key: N, nest; E, entrance to the resting cavity; L, deserted latrines; S, rooting stipe of the fruit body. The folding scale is 51.5 cm long. The photograph is original.

It may be significant that these three observations were accomplished close together in the same geomorphological area, particularly when the breeding of the first two species is compared.

### 3.8 Long-term nesting at the same site

#### Continued nesting

Some species of mammals, such as foxes, are well known to use the same nest or nesting site for long periods, even for two or more generations, while such behaviour in moles was first documented recently by us (Sagara et al., 1993a; Sagrara, 1999, 2009; Sagara and Fukasawa, 2014). For example, *O. mizura* moles nested almost continuously for more than 22 years in one site, from before 1977 to at least 1999 (**Table 3** and **Figure 15**). Here, excavation work was conducted nine times, and the nest was removed seven times, between 1981 and 1989. Despite this disturbance, the moles remained at that site. There are more examples of such persistent use of the same nesting site (Sagara, unpublished).

#### Intermittent nesting

A different case in which the moles nested at the same site with a lapse of about 17 years is shown in **Figure 16** (Sagara, unpublished). This case reflects the repeated intermittent use of a site, since our mushroom watching at that site was conducted annually between 1985 and 2013 (except 1986) but the occurrence of *H. sagarae* was observed only in 1985 and 2002. We observed another similar example over 26 years between 1970 and 1996, where the inhabiting species changed from *M. imaizumii* of Talpini to *U. talpoides* of Urotrichini (Sagara and Okabe, unpublished).

**Figure 16.**
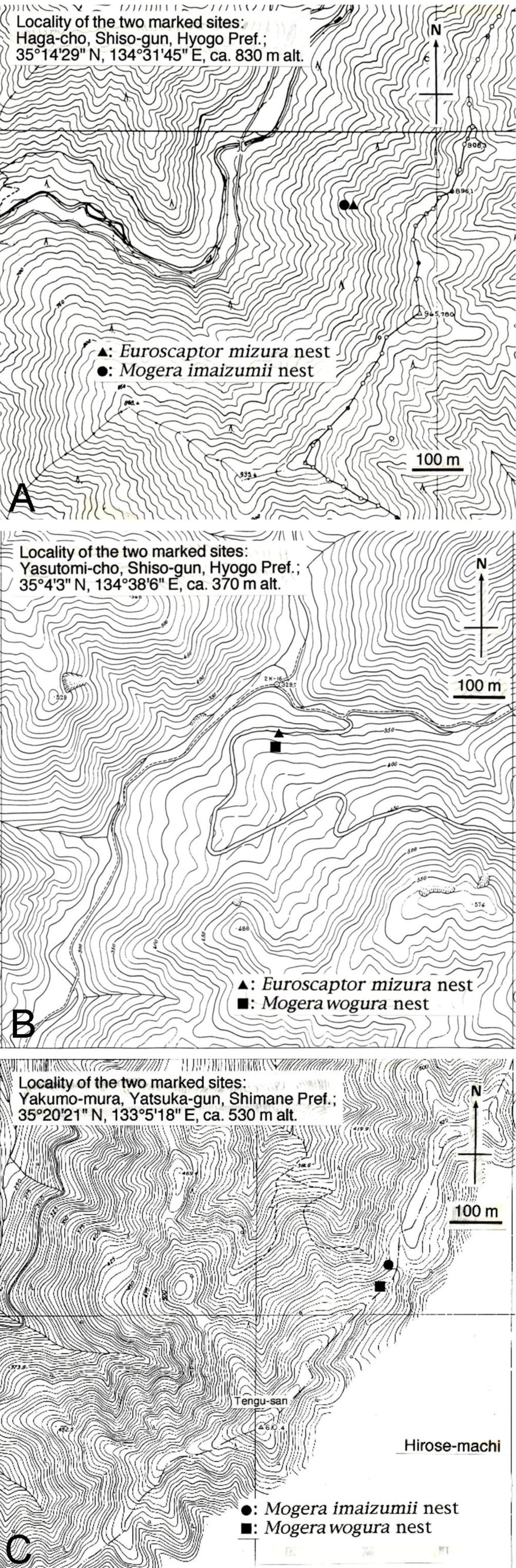
Occurrence of the mushroom *H. sagarae* at the same site in two non-consecutive years, indicating repeated intermittent use of the same nesting site by different individuals. Key: H, *H. sagarae* fruit bodies; F, beech *Fagus crenata* trees. (**A**) Photograph 30 September 1985, showing the first discovery of that mushroom. (**B**) The soil profile at the mushroom fruiting point on 3 October 1985; the nest was located just ahead of this profile but was not removed on this occasion. Key: NL, new latrine. (**C**) Photograph 5 October 2002, showing the second discovery of that mushroom. Key: S, spade. (**D**) The soil profile at the mushroom fruiting point on the same day. Key: DL, deserted latrine; N, nest (abandoned and deteriorated). The thick lines on the folding scale are at 10-cm intervals. This site was used by *M. wogura* on both occasions in 1985 and 2002, as determined by tunnel diameter measurement. Locality: Ashiu, Miyama-cho, Nantan-shi, Kyoto Prefecture, Japan, 685 m alt. Photographs are original.

The behaviour of persisting in a particular nest was observed even during excavation work. The inhabiting mole, which had initially fled from the site when digging began, returned to the site and desperately sought its nest during the silence when the work was suspended. Notably, this behaviour provided an opportunity to capture the mole (Sagara et al., 1989; Sagara et al., 2008b). In addition, when we left the site after digging and removing the nest without replacing the soil, the mole made new mole hills around the empty nest chamber, seeking its own nest (Fig. 25 in Sagara, 1999). Such behaviour gives the impression that moles may have attachment or affection to their own nests and nesting sites.

#### Inhabitant changes

Japanese Talpini individuals live about 4 years at the most (Abe, 1968; Yokohata, 1994). Therefore, the inhabitants during the long-term nesting situations described above must have changed (Sagara et al., 1993a). This assumption was experimentally confirmed using mushroom occurrence as an indicator of mole nesting (Sagara and Fukasawa, 2014). Here, the nest and the nest inhabitant were taken away, and the mushroom refruited later, indicating a different individual had come to nest there. Such inhabitant change occurred not only within the same species but also between different species (**Figure 17**).

**Figure 17.**
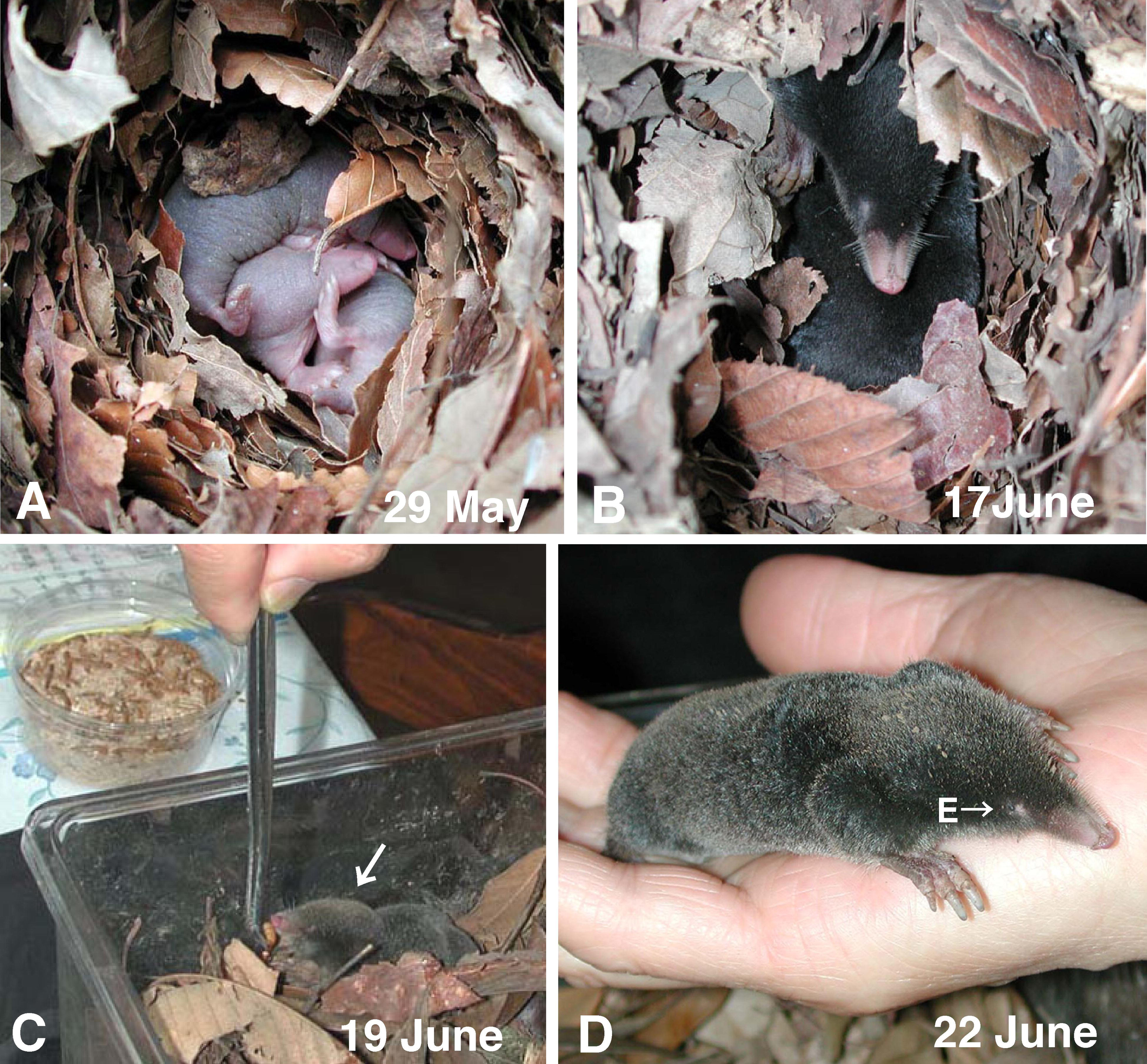
Evidence of the inhabitant change accompanying nest reconstruction at the same site. (**A**) Soil profile with a mole nest (N) on 16 November 1985, obtained by the indication of *H. sagarae* fruiting (H) on 24 September 1984. That nest and its *M. wogura* inhabitant were removed on this occasion. Key: E, entrance to the resting cavity. (**B**) The *M. wogura* individual removed from this location on 15 November 1985 (the day before excavation). (**C**) The soil profile with a reconstructed nest (N) on 7 October 1994, 9 years later. *H. sagarae* (H) refruited on the nest. The new inhabitant was *M. imaizumii*, based on tunnel diameter measurement. Key: L, deserted latrines colonised by mycorrhizas. (**D**) Reference *M. imaizumii* specimen captured near this site on the same occasion as A and B (15 November 1985). Key; S, shrubs (*Symplocos* sp.) to identify the site. The folding scale is 51.5 cm long. The bars in **B** and **D** denote 5 cm. For further information, see Site 3 in Sagara and Fukasawa (2014). Photographs: **A** and **C** from Sagara (1999), reproduced with permission; **B** and **D** original.

## 4. Discussion: moles as forest inhabitants

While studies of mole lives in the forest are scarce compared to those in open areas, all mole nesting examples studied here were from forests in mountains and hills despite undoubtedly occurring in open areas. However, the tripartite association and habitat-cleaning symbiosis among moles, mushrooms, and trees, as schematically shown in **Figure 6**, suggests that their coexistence has a long history (Sagara 1998a). While the following discussion is devoted to understanding moles as forest inhabitants, the distribution of some mole species in mountainous regions is already known (Imaizumi, 1960).

**A.** Godfrey and Crowcroft (1960) wrote on the mole *T. europaea*: ‘it is most abundant in deciduous woodland and those arable areas adjoining it’; ‘there is doubt as to whether it can survive in the soils of mature coniferous forests’. Stone and Gorman (1991) noted that, without explaining, *T. europaea* was originally a deciduous woodland inhabitant. ‘Deciduous woodland’ may be interpreted here as forests of Fagaceae, Betulaceae, and Salicaceae, in the context of mycorrhizal symbiosis (**Section 3.1**).

**B.** Reclamation of the forest for agricultural land by human beings has occurred only recently in Earth’s history. Consequently, moles must have lived in the forest long before this change.

**C.** Moles appear to have originally used fallen broad leaves as nest material and to have established this way of nest making, although they may use grasses and even plastics when available (Gorman and Stone, 1990; Sagara, unpublished). Fallen broad leaves, particularly those of deciduous trees, appear to be best for nest making since the material should: (1) be soft and flexible enough to be carried through the tunnels; (2) be easily manipulated for making the nest wall by laying and pressing the material; (3) provide good absorbency and heat retention.

**D.** Such long-term nesting at the same site as described above may be possible only if the excrement deposited near the nest is soon cleaned away. The proposed ‘habitat-cleaning symbiosis’ (**Section 3.3**) is significant here. Moles do not eat mushrooms or tree roots, but the former must rely on the latter to have their nesting sites cleaned. This relationship may be why moles persist in nesting at sites where that cleaning system has supposedly been established. In this context, we would argue that moles might even know where to nest in relation to mushrooms and plants (Sagara, 1999).

## 5. Future questions

**A.** Are mole nesting sites that should be discovered by the mushroom occurrence really so rare? The studied examples are few (**Table 1**). However, nesting sites located near mushroom occurrence might not be so rare, as suggested by **Figure 11**, which shows the distribution of mole nesting sites in an area that has been relatively intensively surveyed. The rarity of the studied examples may be because mushrooms are easily overlooked. Even when they are detected, they are often not brought to the attention of scientists unless their finder is keen on their ecology.

**B.** The worldwide ranges or alternative species of causal animals, host plants, and cleaner mushrooms in that tripartite association (**Figure 6**) remain unknown. For example, the mushroom *H. radicosum* has been found in North America (Bessette et al., 1997); what would be the causal animal there? Why are cases with mice similar to those mentioned in **Section 2.1** not found in Japan, despite ecologically equivalent mouse species, *Apodemus speciosus* (Temminck) and *Apodemus argenteus* (Temminck), living there alongside moles? Can Pinaceae or other conifers be hosts for *H. radicosum* and *H. sagarae*?

**C.** The biological processes of excrement decomposition in mole latrines should be studied in detail. Based on such studies, the involvement of bacteria and invertebrates in the habitat-cleaning symbiosis will be clarified, and the definition of the association can be extended to a more complex (tetrapartite or pentapartite) relationship.

**D.** How do the mushroom inocula arrive at and colonise the mole latrine? What are the inocula, spores, or hyphae?

**E.** Why does the mushroom *H. sagarae*, and potentially also *H. radicosum*, occur only in wild mammal latrines but not in artificial environments? This mushroom has never been experimentally grown on the forest ground by applying nitrogen compounds to soil, unlike ammonia fungi such as *H. danicum* (Sagara et al., 2008c), while *H. sagarae* is easily grown to fruit in axenic culture (Ohta, 1998). Are *H. sagarae* and *H. radicosum* always mycorrhizal, or can they also be saprotrophic under certain natural conditions?

**F.** The possible variation in mushroom species, as shown by the division of *H. radicosum* into two species (Eberhardt et al., 2020), may need to be carefully studied.

**G.** Why do moles persist in using the same nests and nesting sites? Do they appreciate the presence of mushrooms and ectomycorrhizal plants in the context of habitat-cleaning symbiosis?

**H.** Do *U. talpoides* have unknown nesting behaviours other than nesting underground as commonly observed? (**Section 2.3**).

## 6. Concluding remarks

As the specific epithet of *H. radicosum* and the Japanese names of *H. sagarae* and *H. danicum* (Nagaeno-sugitake and Asinaga-numeri, respectively) indicate, these mushrooms have long-rooting stipes underground. These stipes lead, deep in the soil, to the causal factor of the fruit body occurrence aboveground, i.e., mole or other mammal excretions underground. Similarly, every mushroom species has its own life habits, and its occurrence (fruiting) aboveground or on other substrata is caused by certain factors particular to that species. Consequently, each mushroom’s occurrence reflects what has occurred at a particular location in the past (Sagara, 1975, 1992, 2021; Sagara et al., 2008c). Therefore, mushrooms are storytellers that one can use to explore nature.

A use of mushrooms as storytellers has revealed certain aspects of mole life, as described above. The outstanding contributions from myco-talpology may be the hypothesis of habitat-cleaning symbiosis and the observation of long-term nesting at the same site involving inhabitant changes.

The tripartite association among trees, moles, and mushrooms to create the habitat-cleaning symbiosis occurs in the forest. However, moles nest also in other habitats, and the cleaning processes that may occur in them should be studied, adhering to the question whether such long-term nesting as observed in the forest can also occur.

## Acknowledgements

We, particularly the first author, wishes to repeat expressing his gratitude to those who were already acknowledged in his previous publications for communicating to him their finds of the mushroom occurrence and for assisting him in the field work. Thanks are due to Prof Emer Dr Heinz Clémençon, University of Lausanne, for linguistic discussions on the term ‘myco-talpology’, and to the museums and herbaria mentioned in the text for keeping the collections.

